# Elevated lysosomal mass and enzyme activity in fibroblasts of the Mediterranean mouse *Mus spretus*

**DOI:** 10.1101/2025.02.05.636718

**Authors:** Melissa Sui, Joanne Teh, Kayleigh Fort, Daniel E. Shaw, Peter Sudmant, Tsuyoshi Koide, Jeffrey M. Good, Juan M. Vazquez, Rachel B. Brem

## Abstract

Failures of the lysosome-autophagy system are a hallmark of aging and many disease states. As a consequence, interventions that enhance lysosome function are of keen interest in the context of drug development. Throughout the biomedical literature, evolutionary biologists have found cases in which challenges faced by humans in clinical settings have been resolved by non-model organisms adapting to wild environments. Here, we used a primary cell culture approach to survey lysosomal characteristics in species of the genus *Mus*. We found that fibroblasts from *M. spretus*, a wild Mediterranean mouse, exhibited elevated lysosomal mass and enzyme activity along with reduced activity of β-galactosidase, a classical marker of cellular senescence, compared to those from *M. musculus*, a related species adapted to human-associated environments. We propose that classic laboratory models of lysosome function and senescence may reflect characters that diverge from the phenotypes of wild mice. The *M. spretus* phenotype may ultimately serve as a blueprint for interventions that ameliorate lysosomal dysfunction under conditions of stress and disease.

## Introduction

Many aspects of metazoan health hinge on the ability of the lysosome-autophagy pathway to recycle damaged macromolecules and to direct cell growth decisions (Shin & Zoncu 2020). Indeed, interventions that act broadly to promote health and longevity often require the lysosome-autophagy system (Aman *et al*. 2021; Hansen *et al*. 2018; Bareja *et al*. 2019). More specific mechanisms to boost autophagy are also of potential clinical interest, particularly for treatment of proteinopathies and aging etiologies (Bonam *et al*. 2019; Hansen *et al*. 2018). In practice, whether and how to stimulate proteostasis machinery to advance organismal health remains an open question, and the literature describing such manipulations is in its infancy (Pyo *et al*. 2013; Leiva-Rodríguez *et al*. 2018; Shin *et al*. 2013; Liu *et al*. 2023; Bhuiyan *et al*. 2013; Sami et al. 2002; Bonam et al. 2019; Grabowski & Mistry 2022).

Against a backdrop of decades of work in laboratory systems, ecologists have cataloged stress-and disease-resistance traits in wild genotypes from unusual niches, perhaps most famously in long-lived animal species (Oka *et al*. 2023; Chusyd *et al*. 2021; Finch 2009). Cell-based surveys represent a powerful approach to find and dissect these natural resilience programs. One particularly fruitful discipline has profiled the variation in chemical stress resistance across primary cells from panels of non-model animals, including, in landmark cases, discovery of the underlying mechanisms (Tian *et al*. 2019; Harper *et al*. 2007; Attaallah *et al*. 2020; Sulak *et al*. 2016). Cellular senescence has also been shown to vary across animal species in *in vitro* models (Attaallah *et al*. 2020; Kang *et al*. 2023; Zhao *et al*. 2018; Gomes *et al*. 2011).

In this study, we aimed to leverage species-specific variation in lysosomal markers within the *Mus* genus, to investigate how evolutionary processes have shaped the function of this organelle. Mouse species in this genus shared a common ancestor 7-8 million years ago, and as they radiated across Eurasia, *M. musculus* subspecies came to occupy human-associated niches, whereas other taxa are still found exclusively in the wild (Pagès *et al*. 2015; Smissen & Rowe 2018). Our previous case study (Kang *et al*. 2023) found differences in senescence behaviors, including lysosome markers, across fibroblasts from *M. musculus* subspecies and *M. spretus*, a wild relative that diverged 1-3 million years ago (Morgan *et al*. 2022; Dejager *et al*. 2009). Here, we aimed to build on these observations to gain a deeper understanding of lysosomal programs in non-model mice, focusing on the contrasts between the genotypes of human commensal and laboratory mice and those of their wild relatives.

## Results

### *M. spretus* fibroblasts exhibit reduced lysosomal **β**-galactosidase activity

We previously reported divergence among fibroblasts from several *Mus* genotypes (Kang *et al*. 2023) in characteristics of cellular senescence, a stress-induced program of cell cycle arrest, resistance to apoptosis, and activation of immune signaling (Campisi & d’Adda di Fagagna 2007; Campisi 2005; Hornsby 2002). Most salient from our initial findings were differences between *Mus* fibroblasts in activity of lysosomal β-galactosidase, which is used as a marker of senescence by virtue of its accumulation to detectable levels at suboptimal pH, due to the expanded lysosomal content of senescent cells (Robbins *et al*. 1970; Magalhães & Passos 2018; Dodig *et al*. 2019; Curnock *et al*. 2023).To explore this variation further, we established a panel of primary tail skin fibroblasts from four wild-derived strains of *M. m. musculus* (PWK/PhJ, BLG2/Ms, CHD/Ms, MSM/Ms); two wild-derived strains of *M. m. domesticus* (ManB/NachJ, TUCA/NachJ); an admixed laboratory strain of mostly *M. m. domesticus* origin (C57BL/6J); a wild-derived strain of the steppe mouse *M. spicilegus* (ZRU); and two wild-derived strains of the Mediterranean mouse *M. spretus* (STF/Pas, SFM/Pas). We subjected each culture to 15 Gy of ionizing radiation and incubated for 10 days, which is sufficient to induce DNA damage response, arrested cell growth, and senescence expression programs in fibroblasts from across the genus (Kang *et al*. 2023, Hernandez-Segura *et al*. 2018). In assays of β-galactosidase, fibroblasts of each genotype displayed the anticipated increase in activity following irradiation, compared to their respective control counterparts (Figure 1A-B and Figure S1). Whereas cells from strains of each species behaved similarly, we observed significant differences between species: irradiated *M. spretus* fibroblasts exhibited β-galactosidase signal ∼2-fold lower than did cells from *M. m. musculus* and *M. m. domesticus,* and staining of *M. spicilegus* cells was between the two extremes (Figure 1A-B and Figure S1). To follow up on this difference, we used an alternative approach for senescence induction via the radiomimetic drug neocarzinostatin, which induces senescence in mouse cells (Ito *et al*. 2018; Correia Melo *et al*. 2016, Hernandez-Segura *et al*. 2018). Fibroblasts treated with neocarzinostatin exhibited a pattern mimicking that we had seen under radiation, with the treatment inducing elevated β-galactosidase activity in all cultures, and cells from *M. spretus* staining lower than those of sister taxa (Figure 1C and Figure S1). Furthermore, the same trend was detectable in cells cultured in the absence of DNA damage: fibroblasts from *M. musculus* subspecies exhibited higher β-galactosidase staining than those of *M. spicilegus*, which stained higher than *M. spretus*. The latter species differences from untreated cells were of smaller magnitude than those manifesting after stress, and this dependence on treatment was statistically significant (Figures 1, S1, and S2). Thus, *M. spretus* fibroblasts exhibited dampened β-galactosidase activity relative to other genotypes, with amplification of the effect under stress. Controls ruled out culture passage number as a driver of the variation (Figure S3).

**Figure 1.**
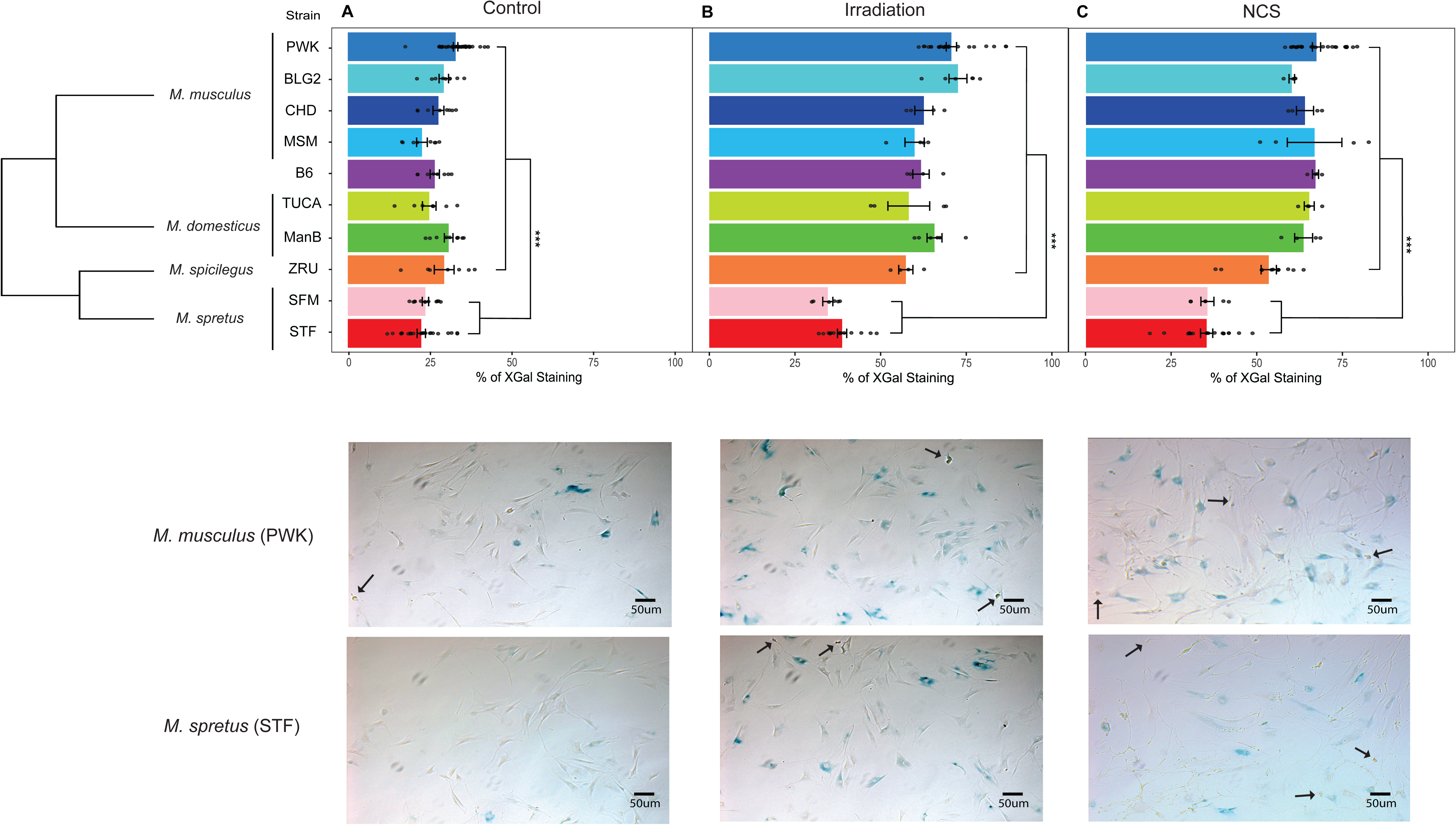
Low β-galactosidase activity in *M. spretus* fibroblasts relative to *M. musculus*. In a given panel, in the plot at the top, each bar length displays the mean percentage of primary fibroblasts of the indicated strain and species that stained positive after administration of the colorimetric β-galactosidase substrate X-Gal. From top to bottom, complete names of the strains are as follows: *M. m. musculus* (PWK/PhJ, BLG2/Ms, CHD/Ms, MSM/Ms), laboratory strain of mostly *M. m. domesticus* origin (C57BL/6J), *M. m. domesticus* (TUCA/NachJ, ManB/NachJ), *M. spicilegus*, and *M. spretus* (SFM, STF/Pas). Representative images of primary fibroblasts from *M. musculus* PWK and *M. spretus* STF are shown at the bottom, with black arrows indicating dead cells. Panels report results from (A) untreated cells, (B) cells treated with ionizing irradiation followed by a 10-day incubation, or (C) cells after 1 hour of neocarzinostatin treatment followed by 24 hours of incubation. Points report biological and technical replicates collected over at least two separate days. Error bars report one standard error above and below the mean. ***, two-tailed Wilcoxon *p* < 10^-7^ comparing *M. spretus* with all other genotypes; *M. spretus* and *M. spicilegus* also differed in each panel (Wilcoxon *p* < 0.05). For (B) and (C), in a comparison between the respective treatment and the untreated control in (A), a two-factor ANOVA with treatment and genotype as factors yielded *p* < 2e^-16^ for the interaction between the two factors. Replicate numbers are provided in Figure S12.

In principle, differences in the senescence-apoptosis fate choice after DNA damage (Childs *et al*. 2014; Zhao *et al*. 2018; Attaallah *et al*. 2020) could contribute to the divergence between mouse genotypes that we had seen in fibroblast β-galactosidase staining. To explore this, we measured caspase 3/5 activity in fibroblasts in response to irradiation and neocarzinostatin treatment (Figure S4). The results showed no consistent patterns of apoptosis either within or between species’ genotypes, arguing against a role for apoptosis activation in species differences in fibroblast β-galactosidase activity.

To establish further the robustness of *Mus* species differences in fibroblast β-galactosidase activity, we considered the dose-response relationship with stress. We compared primary fibroblasts from *M. m. musculus* PWK and *M. spretus* STF as representatives of their respective species, in each case assaying β-galactosidase upon irradiation at increasing doses. The results revealed a consistent ∼2-fold difference between the genotypes at each dose (Figure S5), a magnitude slightly exceeding the species divergence in the untreated control, consistent with our survey across genotypes at fixed stress doses (Figure 1). These data ruled out a switch by *M. spretus* fibroblasts into a high-amplitude, *M. musculus*-like program above a certain threshold of stress exposure. We conclude instead that *M. spretus* cells are hard-wired for lower β-galactosidase activity in all tested conditions—including the resting state.

To advance our search to understand *Mus* species differences in β-galactosidase activity in the fibroblast model, we next focused on regulation of the enzyme itself. Transcriptional profiling data from fibroblasts revealed 1.5 to 2-fold higher expression of the lysosomal β-galactosidase *GLB1* in *M. musculus* fibroblasts than in those of *M. spretus*, regardless of treatment (Figure S6A). Allele-specific expression measurements in stressed and control fibroblasts from an interspecies F1 hybrid made clear that this change was regulated in *cis*: that is, the *M. musculus* allele of *GLB1* drove higher expression of its own encoding locus than the *M. spretus* allele, when both were in the same nucleus (Figure S6B). Thus, the species changes in β-galactosidase activity we had noted in terms of cell biology in fibroblasts (Figure 1) were mirrored by regulation of gene expression, including robust differences between genotypes in unstressed control conditions.

### No consistent divergence in senescence signaling across *Mus* species fibroblasts

We reasoned that the differences in β-galactosidase activity in fibroblasts among the *Mus* species we tested were likely mediated by mechanisms unrelated to senescence itself. As a first investigation of this idea, we explored senescence signaling, using as a readout p21, a key regulator of cell cycle arrest in senescence (Gu *et al*. 2013; Yew *et al*. 2011; Kreis *et al*. 2016). We quantified levels of p21 protein in response to either irradiation or neocarzinostatin, with fibroblasts from *M. m. musculus* PWK and *M. spretus* STF as a testbed. Results showed the expected induction of p21 in fibroblasts from both species in response to irradiation or neocarzinostatin (Figure S7). In investigating quantitative patterns among these p21 induction behaviors, we noted some species-specific differences, though they were not of consistent direction: *M. spretus* cells induced p21 protein abundance more than did *M. musculus* under neocarzinostatin treatment but less following irradiation and in untreated controls (Figure S7). These changes could reflect the differential activity of alternate senescence signaling pathways (Saul *et al*. 2023) between species in some treatments. However, given our focus on lysosomal β-galactosidase differences between *M. musculus* and *M. spretus* across all tested conditions (Figure 1), we concluded that regulation of p21 abundance was not a consistent correlate of this phenotype (although our results do not rule out p21 localization differences between the genotypes). Indeed, in analysis of mRNA expression profiles (Kang *et al*. 2023), we detected no significant difference in induction of the p21 gene (*Cdkn1A*) between *M. musculus* and *M. spretus* fibroblasts in response to stress (Figure S8).

### Increased lysosomal enzyme activity and lysosomal mass in *M. spretus* fibroblasts

We hypothesized that the higher β-galactosidase expression and activity we had seen in *M. musculus* fibroblasts could be a consequence of heightened need owing to failures elsewhere in the proteostasis system, relative to cells of *M. spretus*. To explore this, we first assayed acidic compartments with the LysoTracker stain, again making use of *M. m. musculus* PWK and *M. spretus* STF as a test system. Conforming to our prediction, quantitative microscopy revealed a ∼2.5-fold increase in the cellular area stained by LysoTracker in untreated *M. spretus* fibroblasts compared with *M. musculus* (Figure 2A), and the species difference persisted in cultures induced to senesce with irradiation and neocarzinostatin (Figure 2A). A separate assay design using flow cytometry to quantify LysoTracker staining yielded similar results, with ∼3-fold higher levels in *M. spretus* cells in all conditions (Figure S9). LysoTracker staining was absent from both *M. musculus* and *M. spretus* fibroblasts following treatment with the lysosomal H+-ATPase poison bafilomycin A1, confirming the dependence of this readout on lysosomal acidification (Figure S10).

**Figure 2.**
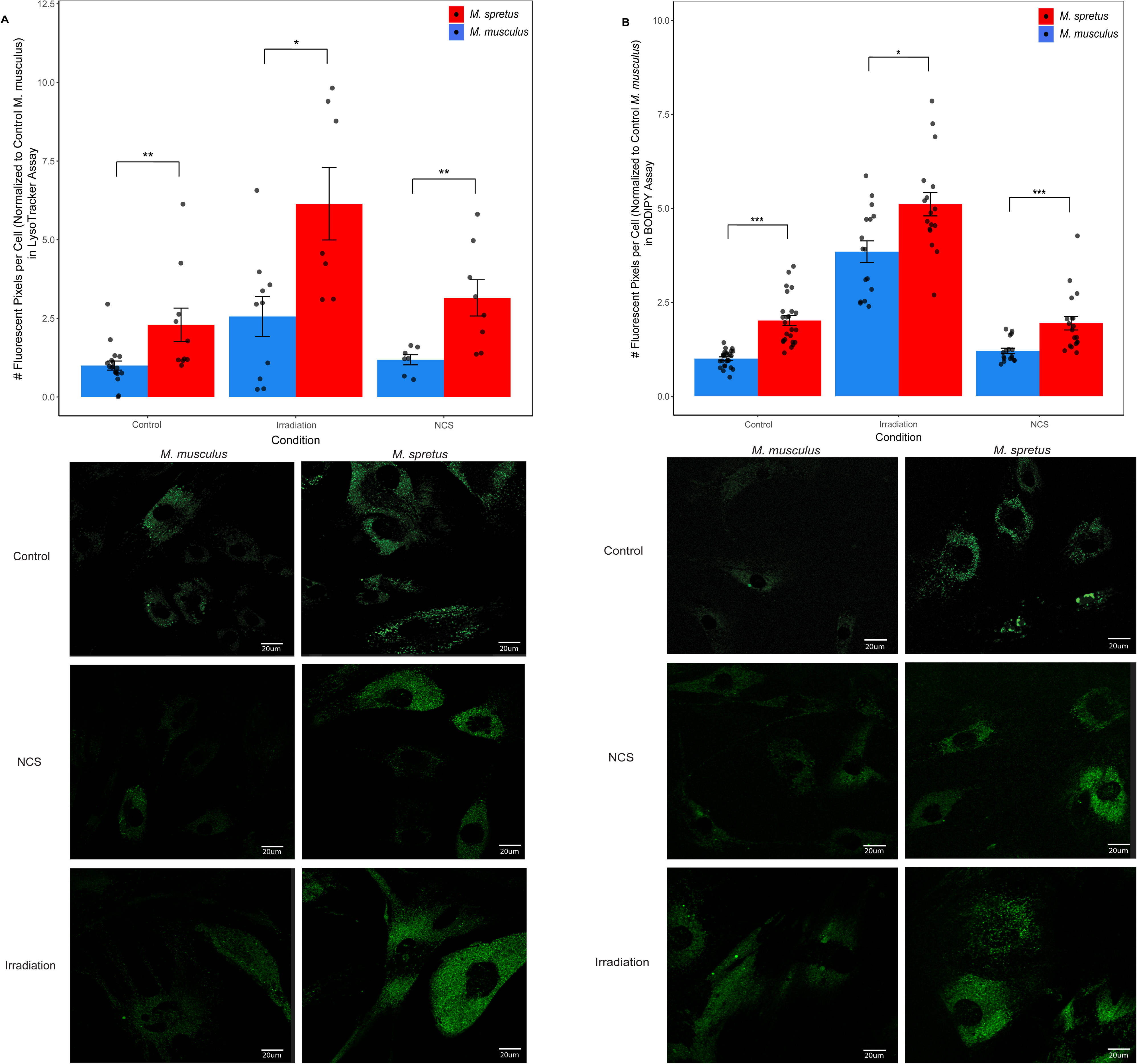
Elevated lysosomal mass and enzyme activity in *M. spretus* fibroblasts relative to *M. musculus*. (A) Shown are results of quantitative microscopy analyses of staining by the lysosomal acidity reporter LysoTracker in primary fibroblasts of the indicated genotype (*M. spretus,* strain STF; *M. m. musculus*, strain PWK). The *y*-axis reports the number of fluorescent pixels per cell, normalized to the control *M. musculus* sample for each experiment. Pairs of bars report results from untreated cells (left), or cells treated with irradiation followed by a 10-day incubation (middle) or after 1 hour of neocarzinostatin (NCS) treatment followed by 24 hours of incubation (right). At the bottom are shown representative fluorescence microscopy images of LysoTracker staining in primary fibroblasts from *M. musculus* PWK and *M. spretus* STF in untreated control, neocarzinostatin (NCS) treatment, and irradiation treatment. (B) Shown are results from assays of cathepsin D activity via BODIPY-pepstatin A fluorescence staining in fibroblasts of the indicated genotype (*M. spretus,* strain STF; *M. m. musculus*, strain PWK). In the plot at the top, the *y*-axis reports the number of fluorescent pixels per cell, normalized to the control *M. musculus* sample for each experiment. Pairs of bars report results from untreated cells (left), or cells treated with irradiation followed by a 10-day incubation (middle) or after 1 hour of neocarzinostatin (NCS) treatment followed by 24 hours of incubation (right). At the bottom are shown representative fluorescence microscopy images of BODIPY-pepstatin A staining in primary fibroblasts from *M. musculus* PWK and *M. spretus* STF in untreated control, neocarzinostatin (NCS) treatment, and irradiation treatment.. Data points correspond to biological and technical replicates collected over at least two different days, and the bar height reports their mean. Error bars report one standard error above and below the mean. *, Wilcoxon *p* < 0.05 , **, Wilcoxon *p* < 0.01, ***, Wilcoxon *p* < 0.001. Replicate numbers are provided in Figure S12.

To investigate further the potential for enhanced lysosomal enzyme function in *M. spretus* cells compared to *M. musculus*, we developed an assay targeting the lysosomal enzyme cathepsin D. This assay employed a BODIPY-labeled pepstatin A conjugate, a fluorescent probe with established use in studies of lysosomal enzyme activity (Boland *et al*. 2008; Lee *et al*. 2010). Measurements from experiments of this design made clear that *M. spretus* fibroblasts exhibited ∼2-fold higher area staining for cathepsin D activity than did *M. musculus* cells under untreated conditions, and similar though slightly dampened differences between the genotypes after irradiation or neocarzinostatin treatment (Figure 2B). Taken together, our results define a trait syndrome in *M. spretus* fibroblasts characterized by elevated lysosomal mass with acidic pH and enzymatic activity, relative to other *Mus* species, under both basal and senescent conditions.

### Lysosomal gene expression divergence between *Mus* fibroblasts

To unearth clues to mechanisms of the divergence between *Mus* species fibroblasts in lysosomal characters, we revisited the expression programs of these cells, this time with a broad survey of lysosome-associated genes. The results revealed marked expression differences between purebred *M. spretus* and *M. musculus* fibroblasts (Figure S11), including high expression in *M. spretus* cells of lysosomal biogenesis factors (namely *Dtnbp1* and members of the *Bloc1s* family); the lysosomal regulator *Lamtor1*; and subunits of the lysosomal H+-ATPase domain *ATP6v1*. Measurements of allele-specific expression in interspecies F1 hybrid fibroblasts established major contributions of *cis*-regulatory variation at many such genes (Figure S11), meaning that the underlying genetic basis of a given gene’s expression change between purebreds could often be ascribed to heritable variants at the locus itself. We conclude that lysosomal gene regulatory changes between the fibroblasts of *Mus* species, controlled in part by *cis*-regulatory variants, parallel the divergence in lysosomal mass and enzyme activity— as expected if *M. spretus* cells made more lysosomes, and populated them to a greater extent with more functional enzymes, as a direct consequence of their distinct regulatory program.

## Discussion

For decades, biomedical researchers have relied on strains from the *M. musculus* clade as the standard for studying lysosomal and senescence biology. In this work, we have demonstrated that lysosomal phenotypes vary quantitatively across *Mus* in fibroblast culture models, and that in contrast to *M. musculus*, cells from the non-commensal mouse *M. spretus* exhibit a program of lysosomal function of the kind that, in the experimental literature, has been associated with healthy aging and disease resistance.

When experimentally induced in laboratory cell models, a backup in the lysosomal-autophagy system triggers compensatory increases in lysosomal mass and number, for which high β-galactosidase activity is a robust marker (Curnock *et al*. 2023; Delfarah *et al*. 2021; Lee *et al*. 2006, Dimri *et al*. 1995, Rovira *et al*. 2022). We propose a similar causal relationship between the phenotypes we have observed as they vary across *Mus* fibroblasts. Under this model, the elevated β-galactosidase activity we have seen in *M. musculus* cells would represent compensation for their limited lysosomal mass and enzyme activity in comparison to *M. spretus* genotypes, even in the absence of stress. The *M. musculus cis*-regulatory allele upregulating β-galactosidase that we have noted in fibroblasts could well represent a genetically encoded component of such a program, a constitutive boost in protein degradation capacity by a regulatory mechanism, in the face of lysosomal defects. Indeed, we consider the broader set of lysosomal genes at which we have detected regulatory variation between the species as compelling candidate components of the mechanism underlying increased lysosomal mass and function in *M. spretus* cells.

Our findings do leave open whether the compromised levels of these phenotypes in *M. musculus* cells represent an ancestral state in *Mus* that was resolved in the *M. spicilegus–M. spretus* lineage, or a derived trait that emerged alongside commensalism in *M. musculus*. In either case, the evidence that we have seen for conservation across *M. musculus* strains and subspecies strongly suggests a history of constraint in this lineage. If the lysosomal behaviors we have studied here prove to have evolved under selection, they could conceivably relate to body size, response to immune challenges, or association with human ecology as they differ between *M. musculus* and *M. spretus* (Dejager *et al*. 2009; Mahieu *et al*. 2006; Kawakami & Yamamura 2008; Staelens *et al*. 2002; Blanchet *et al*. 2011; Pérez del Villar *et al*. 2013; Harr *et al*. 2016).

Ultimately, as its mechanism is revealed, the elevated mass and enzyme activity of lysosomes that we have seen in *M. spretus* fibroblasts are of interest in the search for mimetics that would boost lysosomal function in a human clinical setting. More broadly, our results provide a way forward for the use of wild mouse species as models for lysosomal function and senescence, including future comparisons to the lysosomes of human cells and their phenotypes in disease.

## Methods

Primary tail fibroblasts were extracted as described by Khan and Gasser (2016). Subsequent culture and experiments used complete medium (DMEM, 10% FBS, 1% pen-strep) in T25 flasks at 37°C, 3% O_2_, and 10% CO_2_. Cells were passaged based on confluence using trypsin. Experimental treatments included 15 GY of ionizing irradiation or 3.6 µM neocarzinostatin. Analysis kits used were the Abcam Ltd. Senescence Detection Kit (Cat. #ab65351), ApoTox-Glo™ Triplex Assay (Promega Cat. #G6321), LysoTracker™ Green DND-26 (Thermo Fisher Cat. #L7526), and Pepstatin A BODIPY™ FL Conjugate (Thermo Fisher Cat. #P12271). Primary and secondary antibodies used in western blots were the rabbit monoclonal Anti-p21 antibody (ab188224, Abcam, 1:1000), mouse monoclonal anti-β-tubulin (T9026, Sigma-Aldrich, 1:1000), Goat Anti-Mouse IgG(H+L) Human ads-HRP (Cat#1031-05, Southern Biotech, 1:5000), and Goat Anti-Rabbit IgG(H+L) Human ads-HRP (Cat#4050-05, Southern Biotech, 1:5000). Additional details of methods available in Supplementary Materials.

## Supporting information

Supplemental Materials (Detailed Methods)

## Acknowledgements

The authors thank Vera Gorbunova, Emilie Tu, Helen Bateup, Samantha Jackson, Linda Wilbrecht, Polly Campbell, and Michael Nachman for animal material; Diana Bautista, Britt Glaunsinger, José Pablo Vázquez-Medina, and Mary West for their generosity with space and resources; Juliana Valencia Lesmes, Mara Baylis, Sam Rider, Harriet Song and William D’Angelo for technical contributions; and members of the Brem lab for helpful discussions. This work was supported by National Institutes of Health R01NS116992 to RBB and R01GM120430 to RBB and JMG. JMV was supported by the National Science Foundation Postdoctoral Research Fellowship in Biology (#2109915) and by the National Institute for Health T32AG000266 and 1K99AG088361.

## Data Accessibility Statement

Data used in this manuscript can be found in an external data repository on Figshare at https://doi.org/10.6084/m9.figshare.c.8086870.v1. RNA-sequencing data used in this manuscript can be found in the NCBI Gene Expression Omnibus (GEO; https://www.ncbi.nlm.nih.gov/geo/) under accession number GSE201217.

## Supplementary figure captions

**Figure S1.**
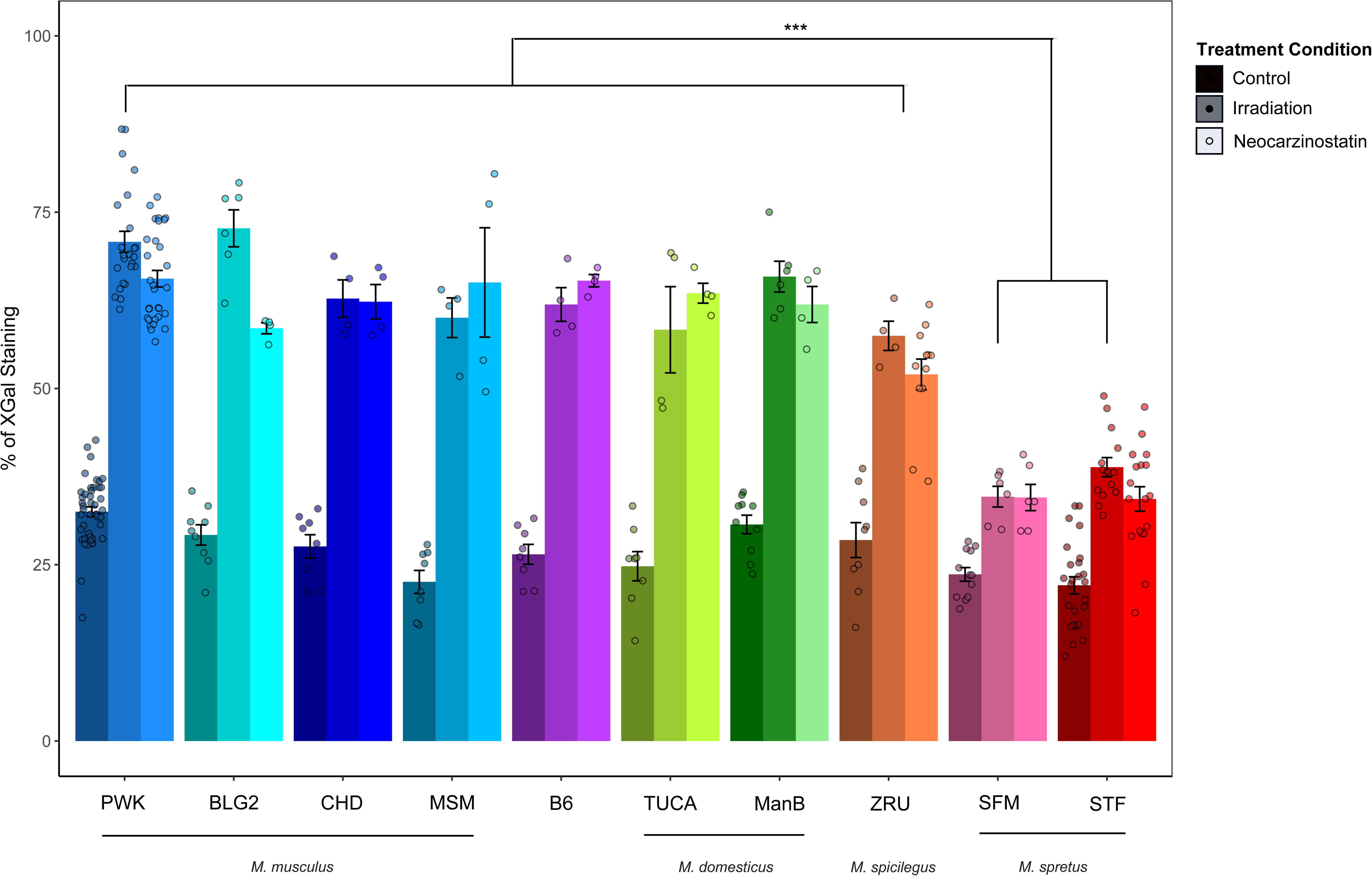
Impact of genotype and treatment on β-galactosidase activity in *Mus* fibroblasts. Data are those of Figure 1 of the main text, grouped by genotype to emphasize treatment effects. A given set of three bars represents results from primary fibroblasts of the indicated strain from three conditions: untreated cells (darkest color), cells exposed to ionizing irradiation followed by a 10-day incubation (medium color), and cells treated with neocarzinostatin for 1 hour followed by 24 hours of incubation (lightest color).

**Figure S2.**
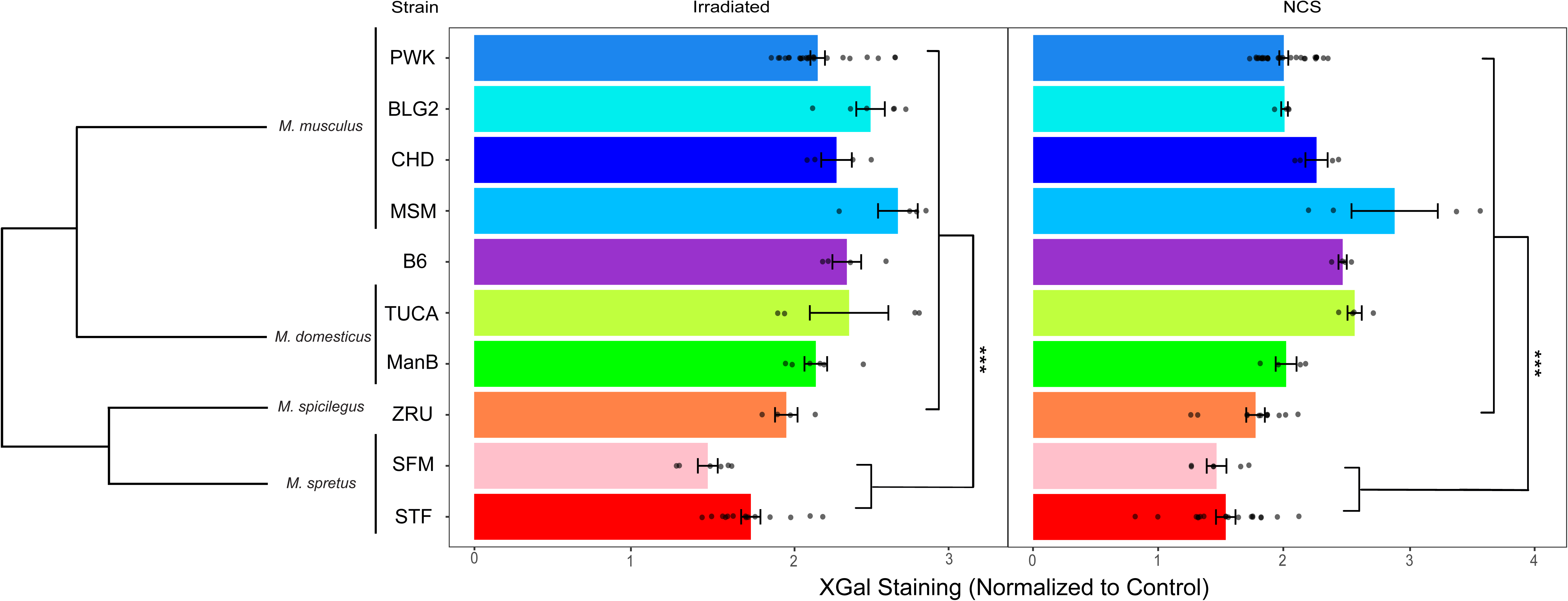
Low normalized β-galactosidase activity in *M. spretus* fibroblasts. Data are as in Figure 1 of the main text, except that measurements from the culture of a given genotype and treatment were normalized to the average of all measurements from untreated controls of that genotype. ***, two-tailed Wilcoxon *p* < 0.001 comparing *M. spretus* with all other genotypes for both treatments.

**Figure S3.**
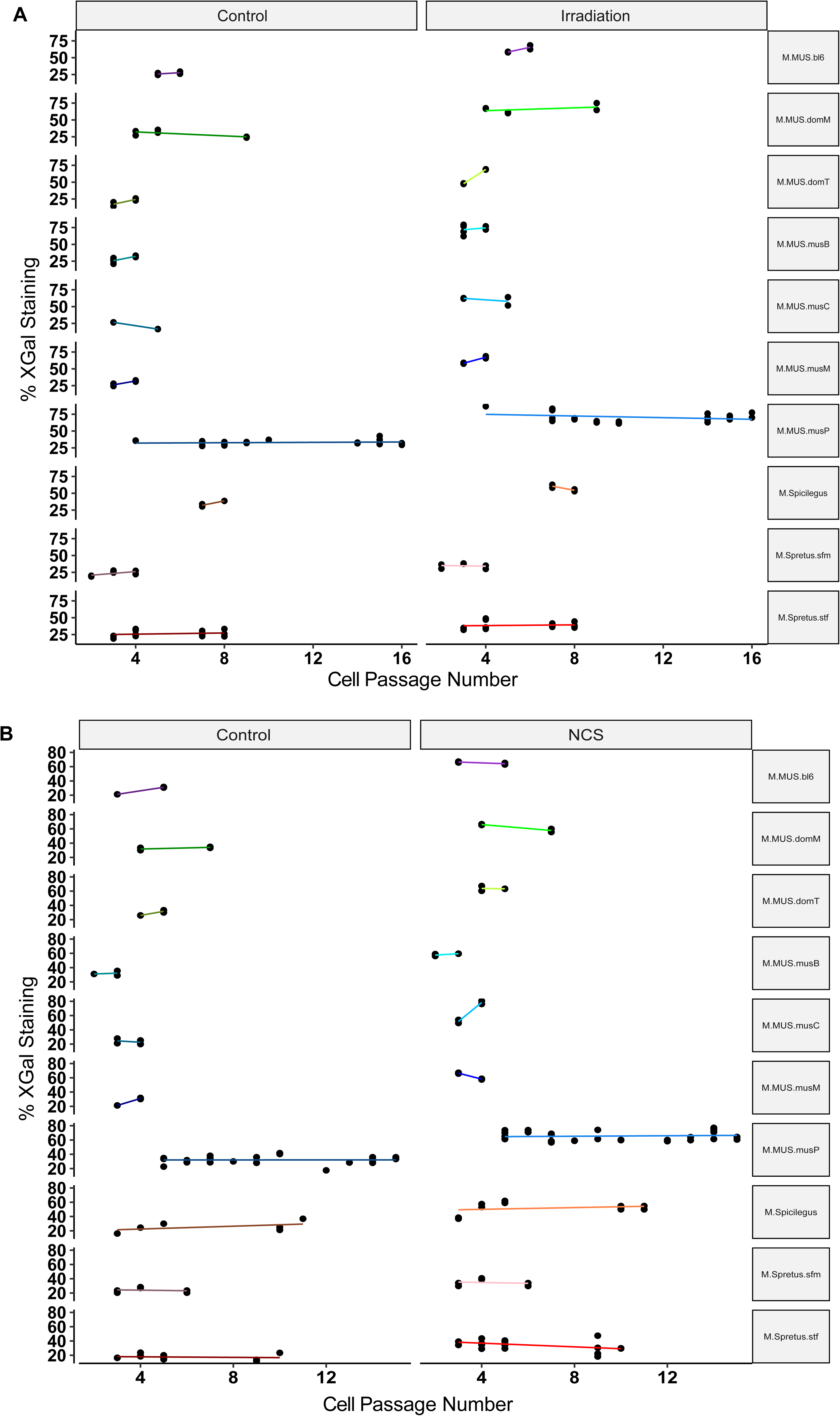
Fibroblast passage number has no detectable effect on X-Gal staining. In a given panel, each row reports results from assays of the β-galactosidase substrate X-Gal on fibroblasts of the indicated genotype shown in Figure 1 of the main text, and each point reports one replicate culture. The x-axis reports the passage number of the respective culture and the *y*-axis reports X-Gal staining. (A) Cells induced to senesce with irradiation (right) and their paired controls (left). (B) Cells induced to senesce with neocarzinostatin (right) and their paired controls (left). Replicate numbers are provided in Figure S12.

**Figure S4.**
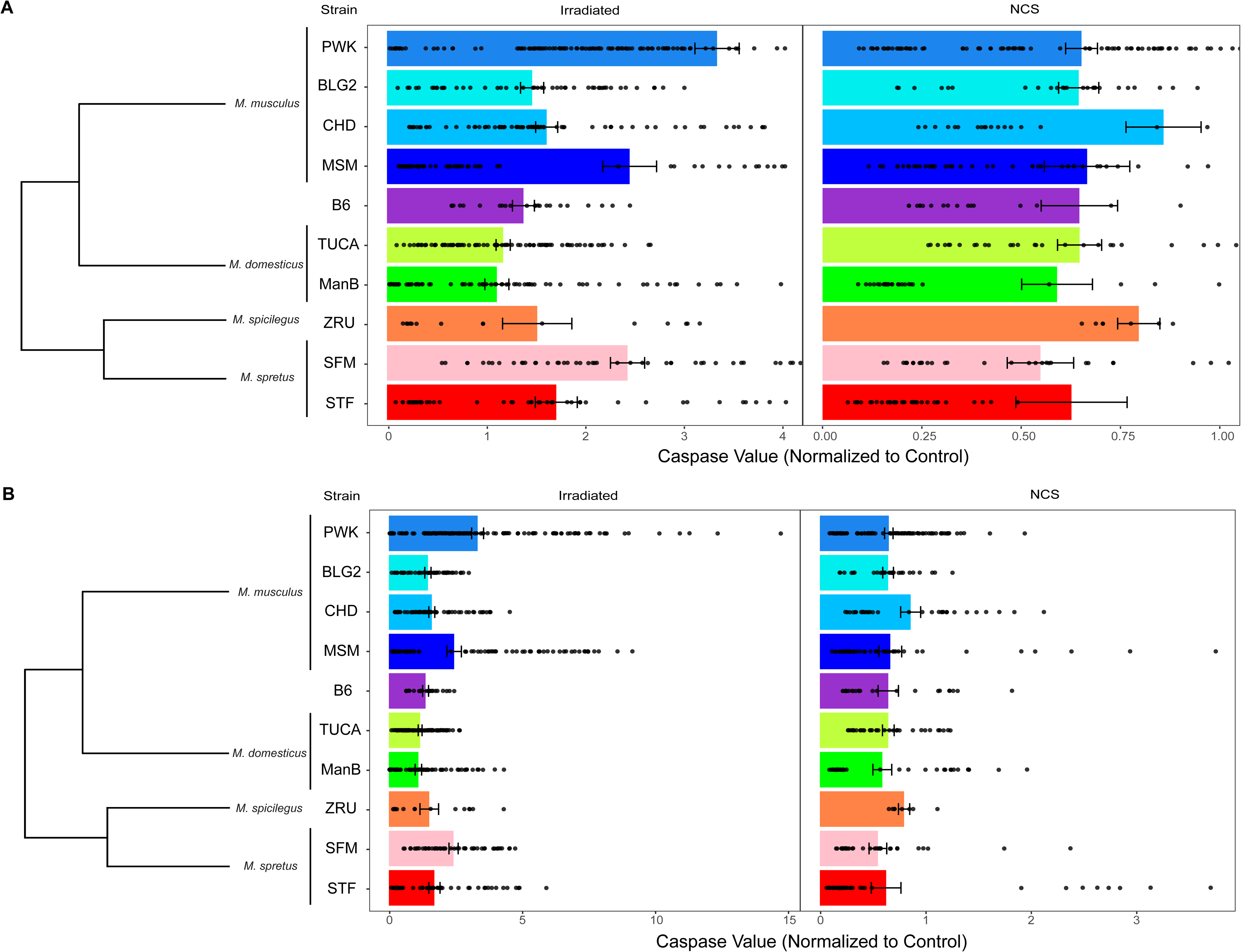
No consistent variation between *Mus* species fibroblasts in apoptosis activity. Data are as in Figure 1 of the main text, except that for a given genotype and treatment, the *x*-axis reports caspase activity of the culture measured with the Apotox-Glo assay, normalized to the average from untreated cultures of the respective genotype for each experiment. (A) Outliers trimmed for visualization. (B) Full data set view. Replicate numbers are provided in Figure S12.

**Figure S5.**
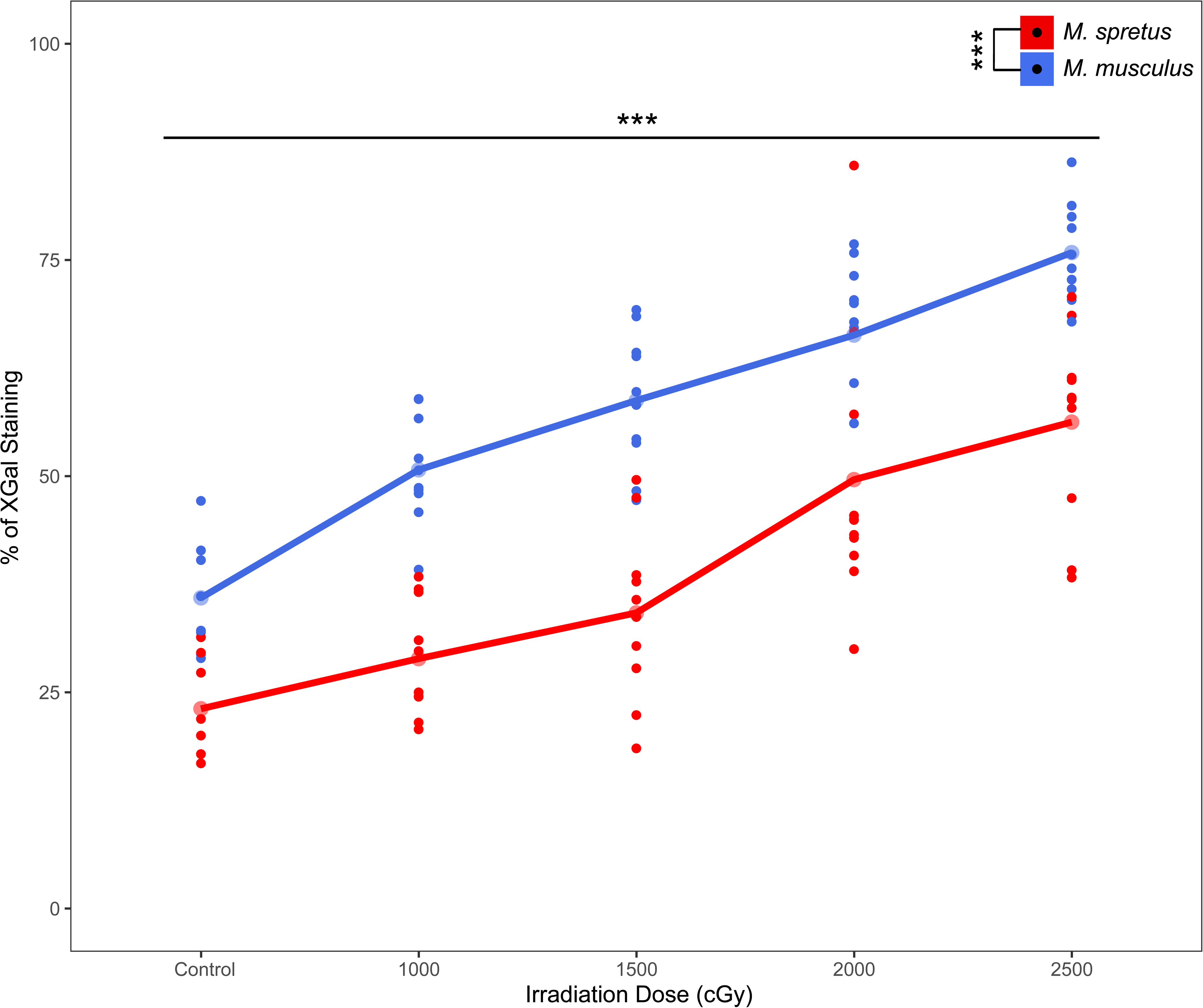
Dose dependence of β-galactosidase staining in *Mus* fibroblasts. Each trace shows results from assays of the β-galactosidase substrate X-Gal on primary fibroblasts of the indicated genotype (*M. spretus*, strain STF; *M. m. musculus*, strain PWK). For a given trace, the x-axis reports irradiation dosage (in centigray, cGy) at 0 (Control), 1000, 1500, 2000, and 2500 cGy, and the y-axis reports the percentage of X-Gal positive cells in untreated controls or in irradiated cells after 10 days of incubation. Smaller points represent biological and technical replicates, and larger points indicate their mean values. ***, two-factor ANOVA with dosage and species as factors yielded *p* < 2e^-16^ for each factor. Replicate numbers are provided in Figure S12.

**Figure S6.**
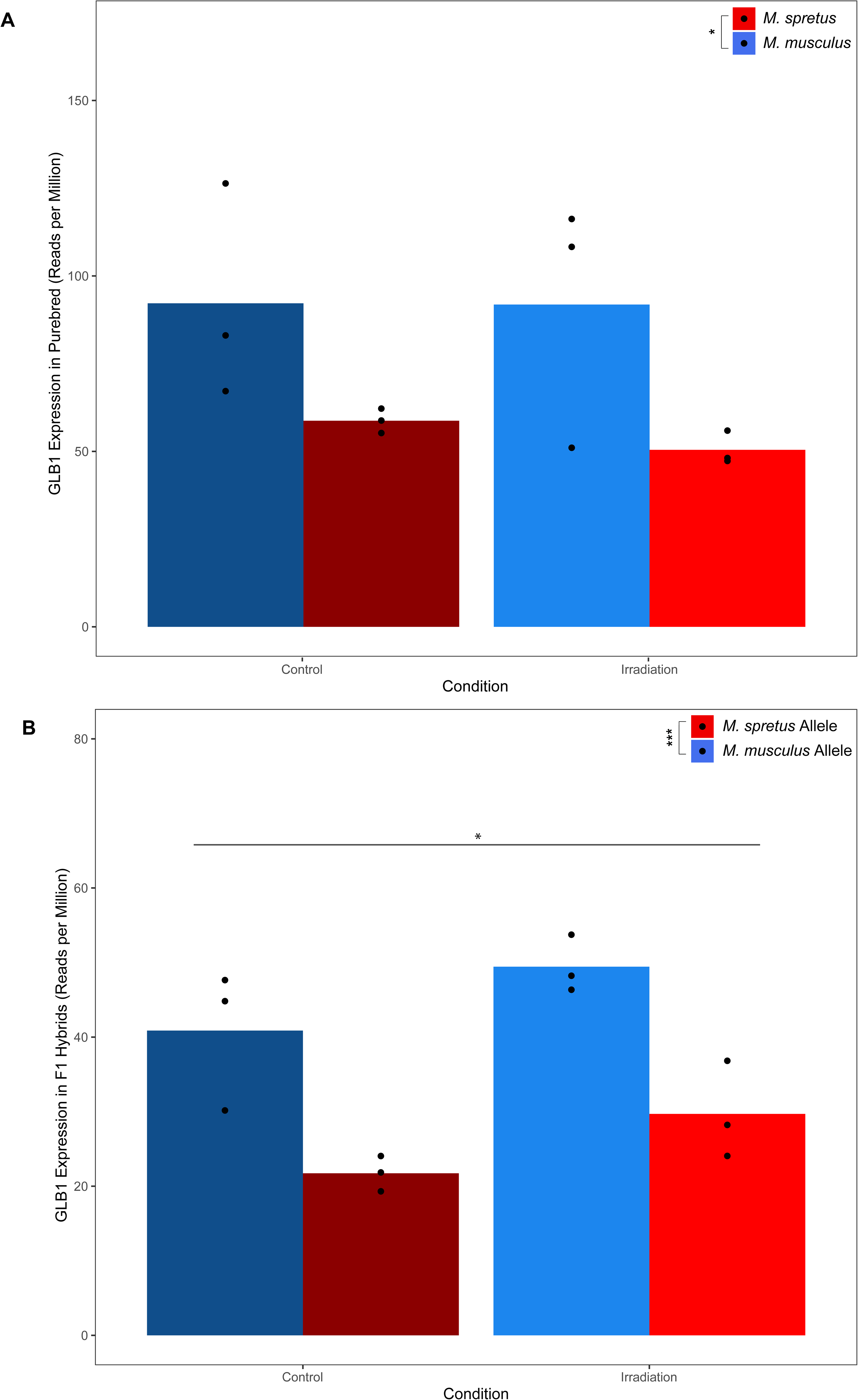
Variation between *Mus* fibroblasts in Glb1 mRNA expression. (A) Shown are measurements of mRNA expression of the β-galactosidase gene from profiling of fibroblasts of the indicated genotype (*M. spretus*, strain STF; *M. m. musculus*, strain PWK), untreated or treated with irradiation followed by a 10-day incubation (Kang *et al*. 2023). Data points within each column represent biological and technical replicates and the bar height reports their mean. Only the effect of genotype was significant in a two-factor ANOVA with condition and species as factors (*, *p* < 0.05). Wilcoxon test yielded no significant difference between the species for each condition. (B) Data are as in (A), except that allele-specific mRNA expression of the β-galactosidase gene was measured from profiling of fibroblasts of an F1 hybrid between *M. spretus*, strain STF and *M. m. musculus*, strain PWK (Kang *et al*. 2023). In a two-factor ANOVA with condition and genotype as factors, both had significant effects (species *p* < 0.0005, ***; condition *p* < 0.05, *). Replicate numbers are provided in Figure S12.

**Figure S7.**
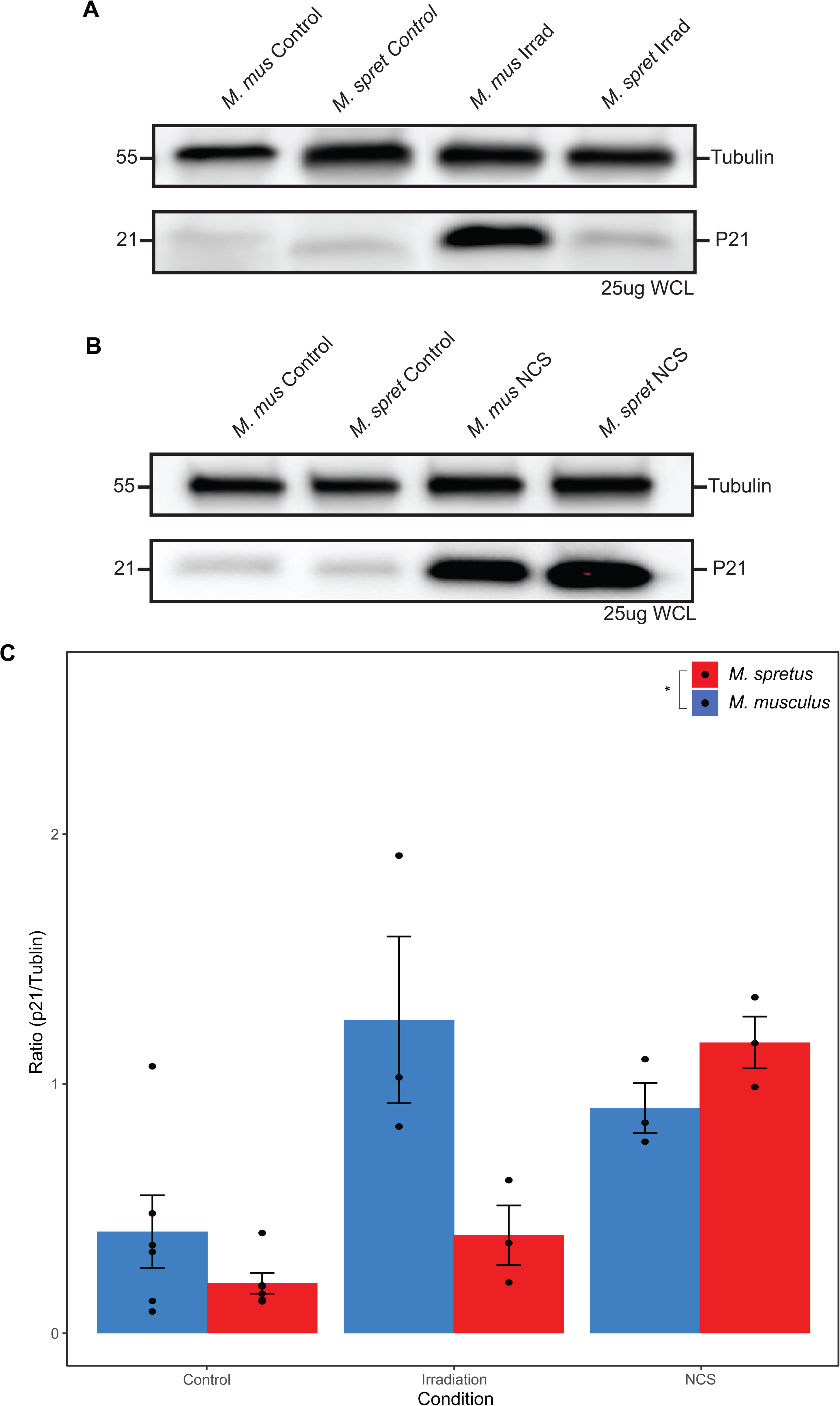
No consistent variation between *Mus* fibroblasts in p21 abundance. Shown are results of Western blot assays of abundance of the senescence regulator p21 in primary fibroblasts of the indicated genotype (*M. spretus*, strain STF; *M. m. musculus*, strain PWK). (A) Top, representative blot showing tubulin (55 kDa) and p21 (21 kDa) levels in 25 µg of whole cell lysate (WCL) from untreated control cells or those irradiated (Xray) and incubated for 10 days to establish senescence. Bottom, blot as at top except that cells were treated with 1 hour of neocarzinostatin (NCS) and incubated for 24 hours to establish senescence. (B) Each column reports quantification of p21 protein abundance normalized to tubulin for the indicated genotype and treatment. Data points correspond to biological and technical replicates collected over at least two different days, and the bar height reports their mean. Error bars report one standard error above and below the mean. No comparisons were significant in pairwise Wilcoxon tests. *, two-factor ANOVA with condition and species as factors yielded p < 0.05 for species. Replicate numbers are provided in Figure S12.

**Figure S8.**
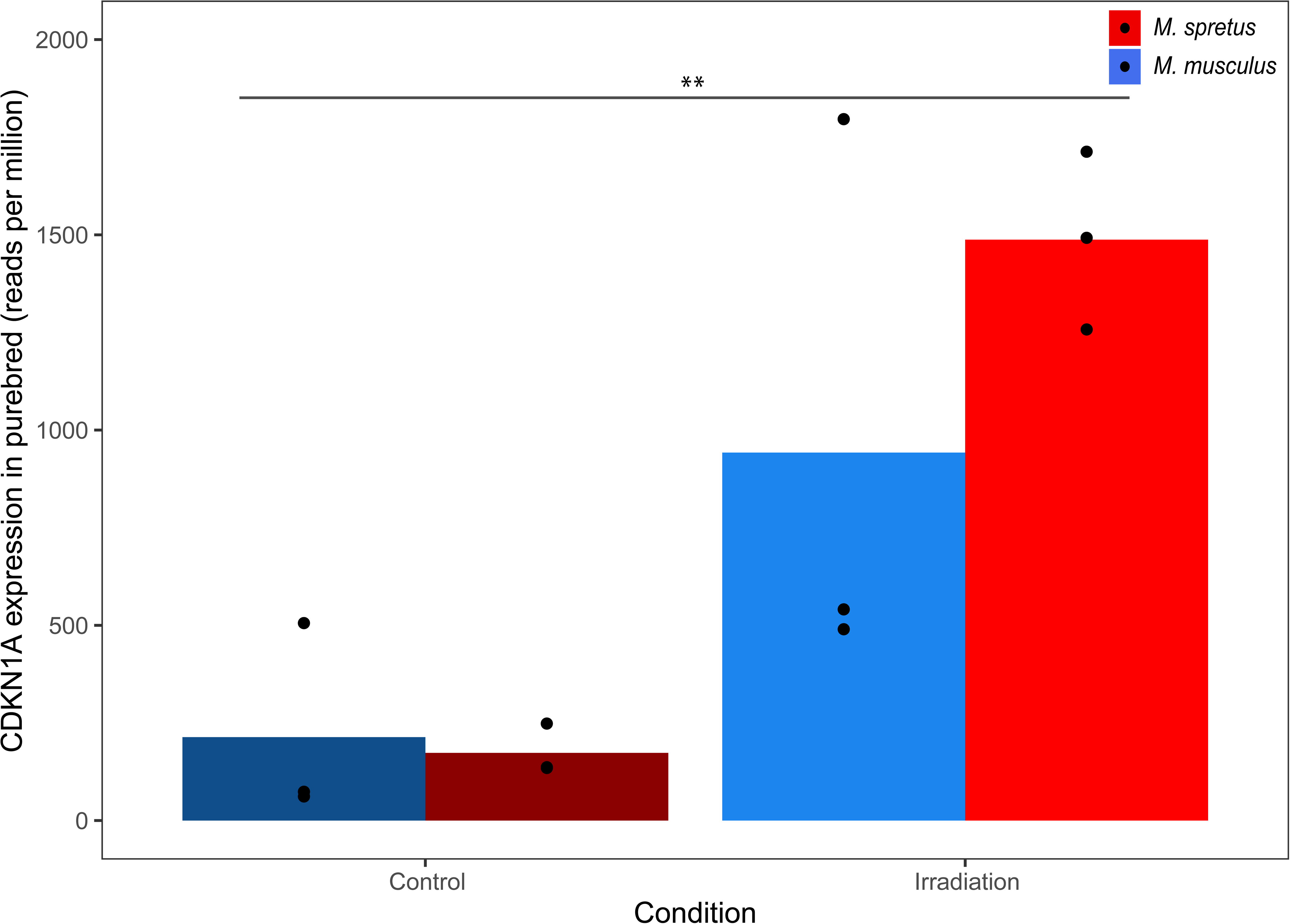
No significant variation between *Mus* fibroblasts in Cdkn1a mRNA expression. Shown are measurements of mRNA expression of the p21 gene Cdkn1a, from profiling of fibroblasts of the indicated genotype (*M. spretus*, strain STF; *M. m. musculus*, strain PWK), untreated or treated with irradiation followed by a 10-day incubation (Kang *et al*. 2023). Data points within each column represent biological and technical replicates and the bar height reports their mean. Error bars report one standard error above and below the mean. Only condition had a significant impact on expression in a two-factor ANOVA with treatment and genotype as factors (p < 0.005, **). Replicate numbers are provided in Figure S12.

**Figure S9.**
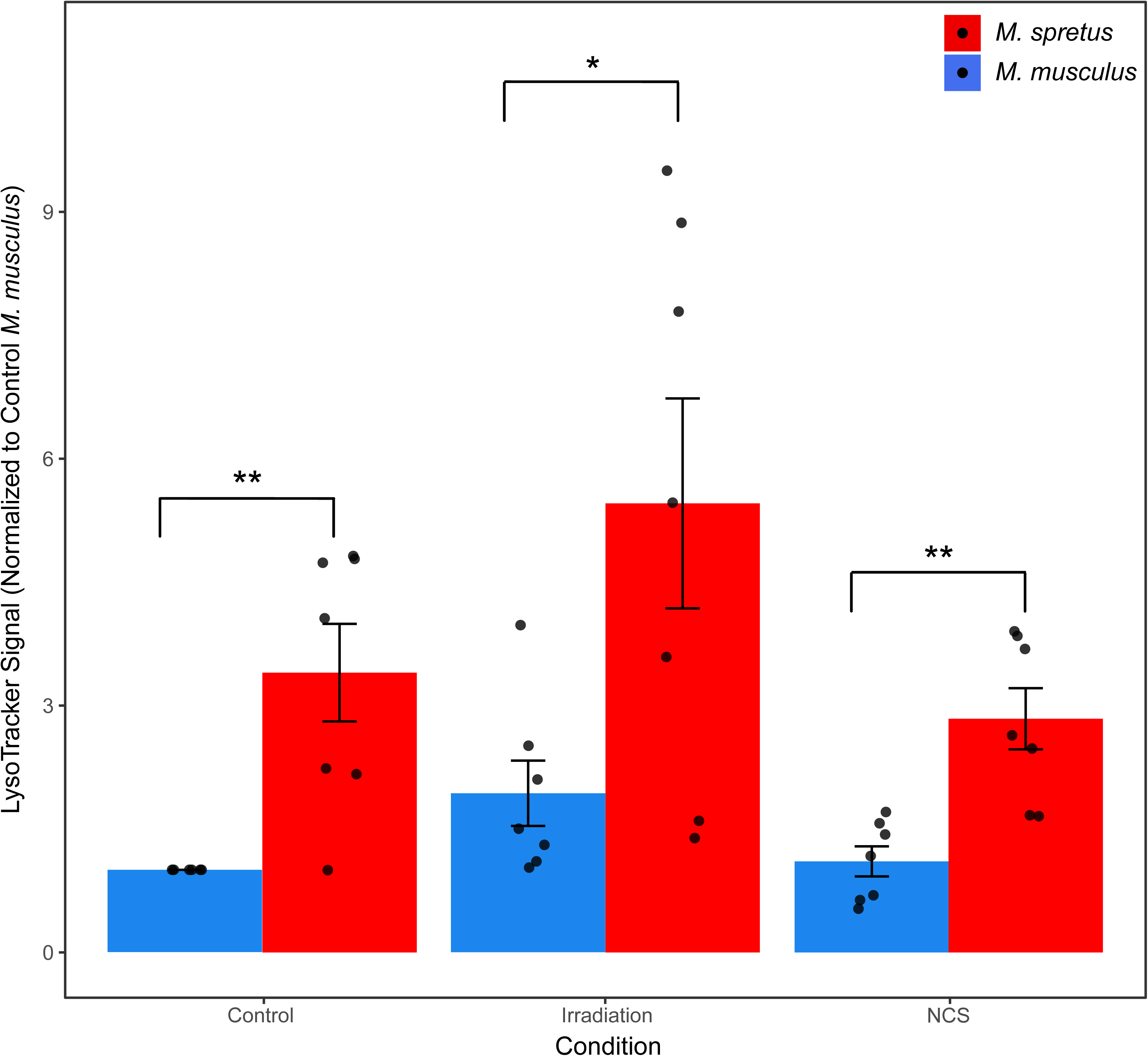
Fluorescent signal quantification of LysoTracker staining indicates enhanced intracellular acidity in *M. spretus* fibroblasts. Shown are results from assays of the intracellular acidity reporter LysoTracker on primary fibroblasts of the indicated genotype (*M. spretus,* strain STF; *M. m. musculus*, strain PWK). The *y*-axis reports mean fluorescence of the indicated culture normalized to the control *M. musculus* sample for each experiment. Data points correspond to biological and technical replicates collected over at least two different days, and the bar height reports their mean. Error bars report one standard error above and below the mean. *, Wilcoxon *p* < 0.05 , **, Wilcoxon *p* < 0.01. Replicate numbers are provided in Figure S12.

**Figure S10.**
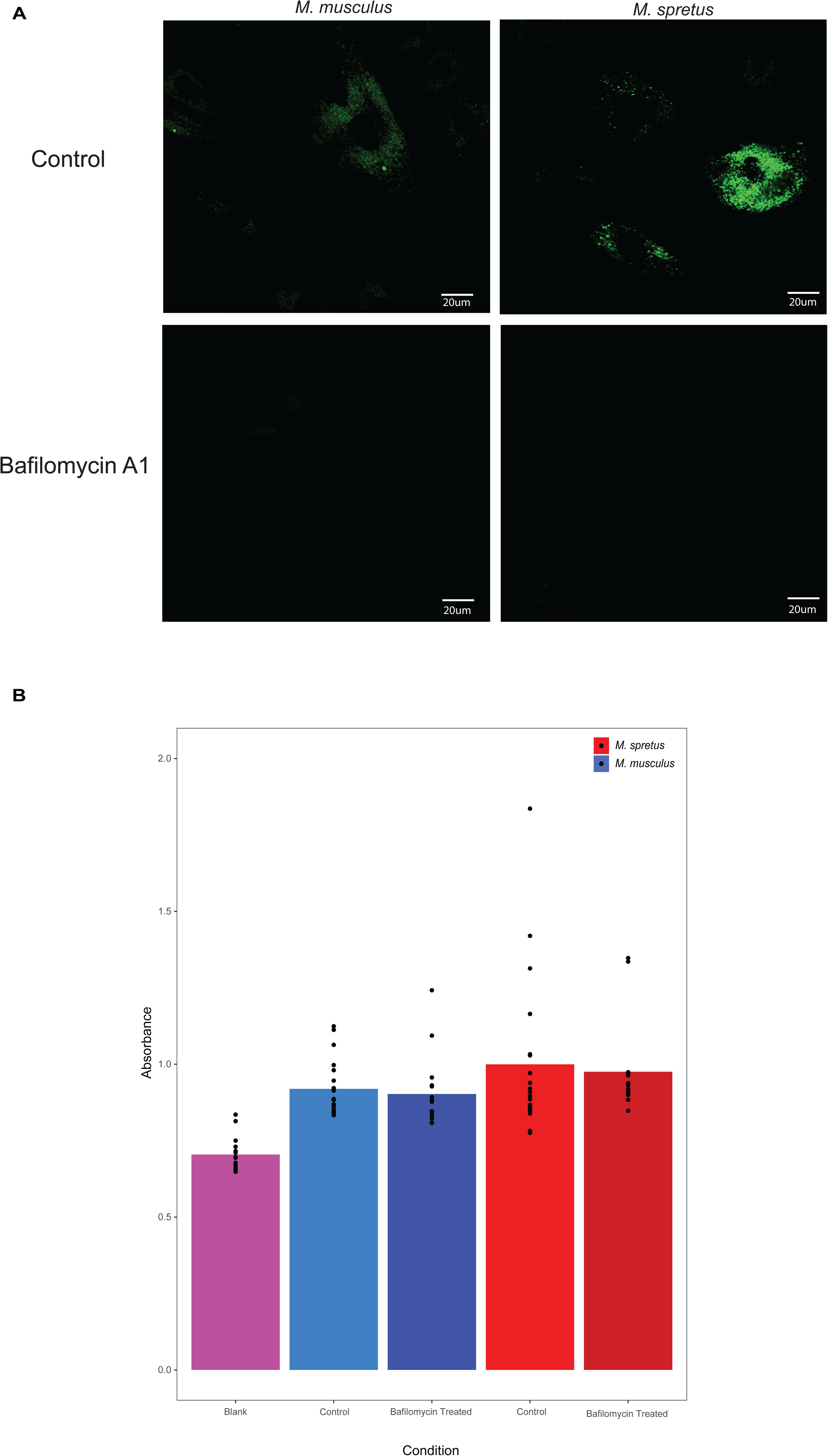
Bafilomycin A1 treatment abrogates LysoTracker fluorescent staining. (A) Shown are representative microscopy images of intracellular acidity reporter LysoTracker in untreated primary fibroblasts of the indicated genotype (*M. spretus,* strain STF; *M. m. musculus*, strain PWK) after bafilomycin A1 treatment. (B) Alamar Blue cell viability stain results are shown for primary fibroblasts of the indicated genotypes (*M. spretus*, strain STF; *M. m. musculus*, strain PWK) either with or without bafilomycin A1 treatment, or a no-cell control (Blank). The y-axis indicates absorbance of cell cultures at 570 nm, corresponding to Alamar Blue after reduction by active cellular metabolism. Data points within each column represent technical replicates and the bar height reports their mean.

**Figure S11.**
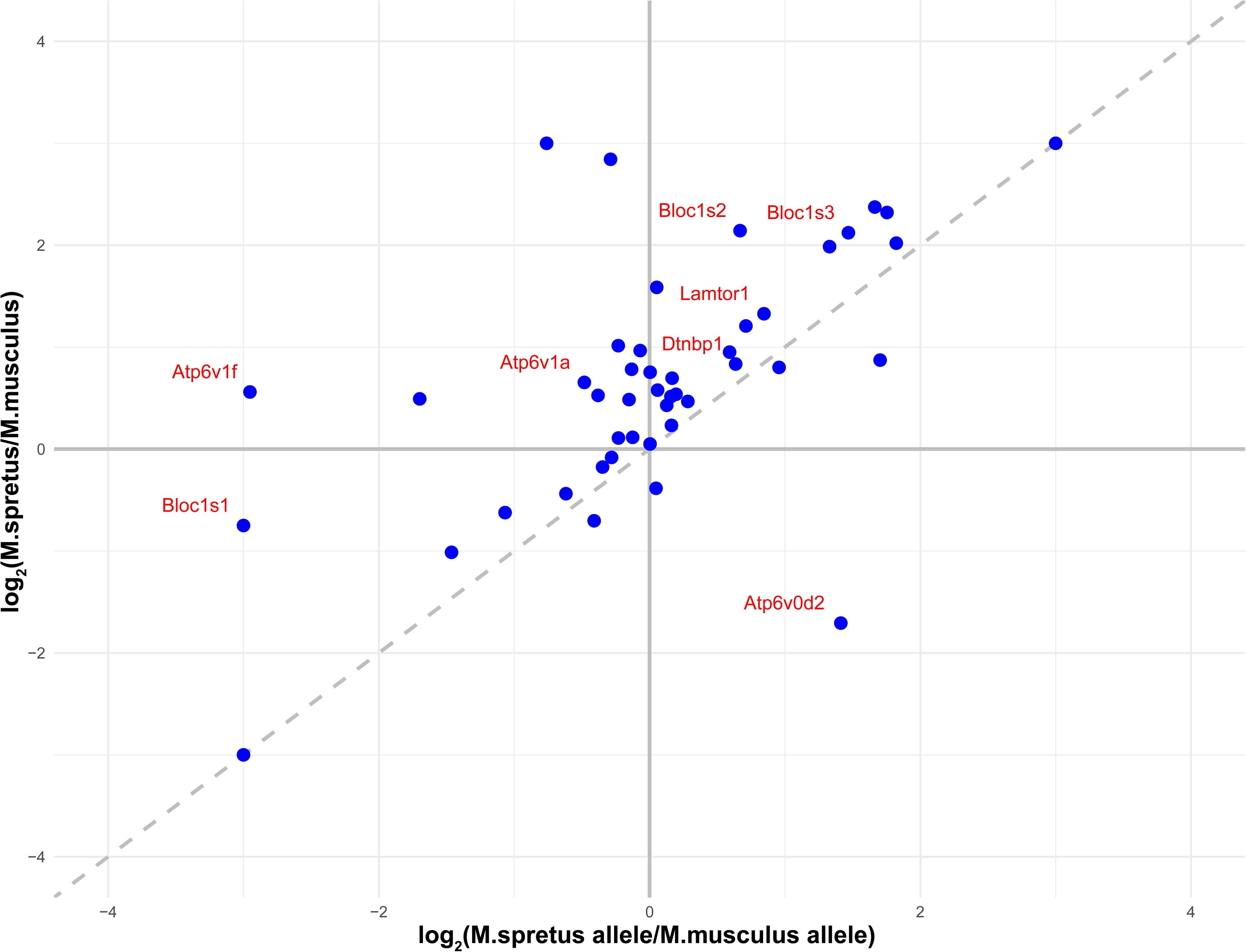
Variation between species fibroblasts in lysosomal gene expression. Each point reports the effect of genetic variation between *M. spretus* STF and *M. m. musculus* PWK on the expression of a lysosomal gene in primary skin fibroblasts from (Kang *et al*. 2023). The *x*-axis shows the ratio of expression of the *M. spretus* allele relative to that of the *M. musculus* allele in an F1 hybrid background, indicating the contribution of *cis*-acting regulatory variants. The *y*-axis displays gene expression in purebred *M. spretus* fibroblasts normalized to that of *M. musculus*. Genes positioned near the diagonal line exhibit expression differences primarily driven by *cis*-regulatory variation. Red-labeled genes are highlighted in the main text.

**Figure S12.**
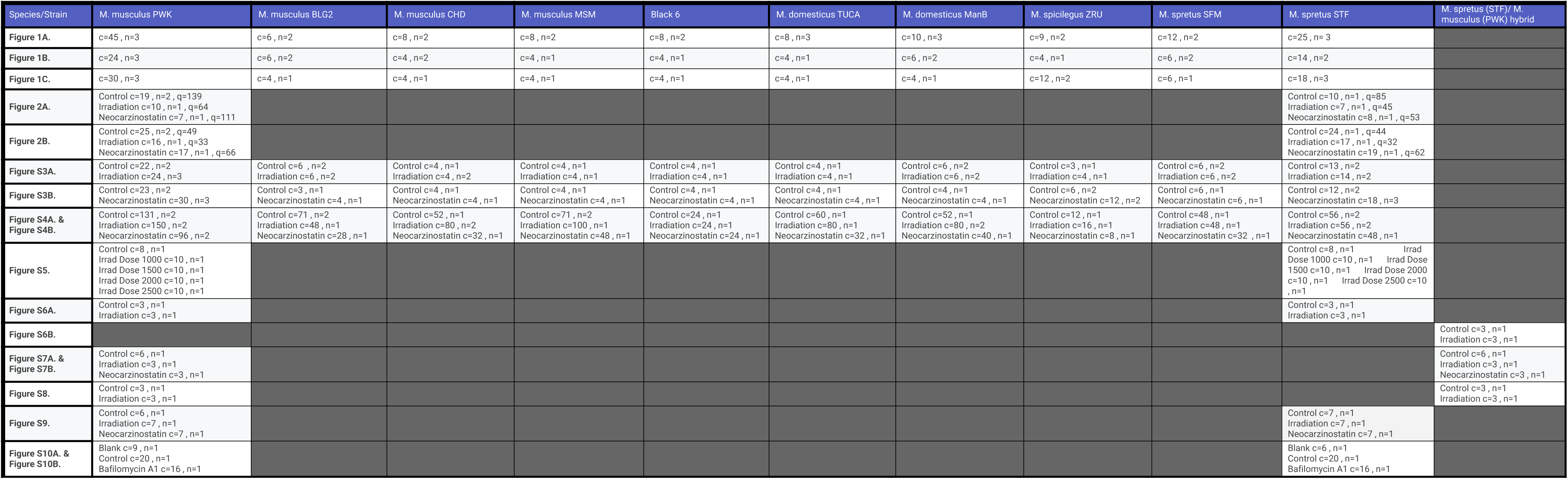
Sample sizes. Shown are sample sizes for each experiment presented in the main and supplementary figures. (n) denotes the number of independent animals used per experiment, (c) denotes the number of independent culture replicates used per experiment, and (q) denotes the number of cells quantified in the experiment.

## References

1. Aman Y, Schmauck-Medina T, Hansen M, Morimoto RI, Simon AK, Bjedov I, Palikaras K, Simonsen A, Johansen T, Tavernarakis N, Rubinsztein DC, Partridge L, Kroemer G, Labbadia J & Fang EF (2021) Autophagy in healthy aging and disease. Nat Aging 1, 634–650.

2. Attaallah A, Lenzi M, Marchionni S, Bincoletto G, Cocchi V, Croco E, Hrelia P, Hrelia S, Sell C & Lorenzini A (2020) A pro longevity role for cellular senescence. Geroscience 42, 867–879.

3. Bareja A, Lee DE & White JP (2019) Maximizing Longevity and Healthspan: Multiple Approaches All Converging on Autophagy. Front Cell Dev Biol 7, 183.

4. Bhuiyan MS, Pattison JS, Osinska H, James J, Gulick J, McLendon PM, Hill JA, Sadoshima J & Robbins J (2013) Enhanced autophagy ameliorates cardiac proteinopathy. J Clin Invest 123, 5284–5297.

5. Blanchet C, Jaubert J, Carniel E, Fayolle C, Milon G, Szatanik M, Panthier J-J & X Montagutelli (2011) Mus spretus SEG/Pas mice resist virulent Yersinia pestis, under multigenic control. Genes Immun 12, 23–30.

6. Boland B, Kumar A, Lee S, Platt FM, Wegiel J, Yu WH & Nixon RA (2008) Autophagy Induction and Autophagosome Clearance in Neurons: Relationship to Autophagic Pathology in Alzheimer’s Disease. J Neurosci 28, 6926–6937.

7. Bonam SR, Wang F & Muller S (2019) Lysosomes as a therapeutic target. Nat Rev Drug Discov 18, 923–948.

8. Campisi J (2005) Senescent cells, tumor suppression, and organismal aging: good citizens, bad neighbors. Cell 120, 513–522.

9. Campisi J & d’Adda di Fagagna F (2007) Cellular senescence: when bad things happen to good cells. Nat Rev Mol Cell Biol 8, 729–740.

10. Childs BG, Baker DJ, Kirkland JL, Campisi J & van Deursen JM (2014) Senescence and apoptosis: dueling or complementary cell fates? EMBO Rep 15, 1139–1153.

11. Chusyd DE, Ackermans NL, Austad SN, Hof PR, Mielke MM, Sherwood CC & Allison DB (2021) Aging: What We Can Learn From Elephants. Front Aging 2, 726714.

12. Correia Melo C, Marques FD, Anderson R, Hewitt G, Hewitt R, Cole J, Carroll BM, Miwa S, Birch J, Merz A, Rushton MD, Charles M, Jurk D, Tait SW, Czapiewski R, Greaves L, Nelson G, Bohlooly-Y M, Rodriguez-Cuenca S, Vidal-Puig A, Mann D, Saretzki G, Quarato G, Green DR, Adams PD, Von Zglinicki T, Korolchuk VI & Passos JF (2016) Mitochondria are required for pro-ageing features of the senescent phenotype. The EMBO Journal 35, 724–742.

13. Curnock R, Yalci K, Palmfeldt J, Jäättelä M, Liu B & Carroll B (2023) TFEBLdependent lysosome biogenesis is required for senescence. EMBO J 42, e111241.

14. De Magalhães JP & Passos JF (2018) Stress, cell senescence and organismal ageing. Mechanisms of Ageing and Development 170, 2–9.

15. Del Villar LP, Vicente B, Galindo-Villardón P, Castellanos A, Pérez-Losada J & Muro A (2013) Schistosoma mansoni experimental infection in Mus spretus (SPRET/EiJ strain) mice. Parasite, 20, 27.

16. Dejager L, Libert C & Montagutelli X (2009) Thirty years of Mus spretus: a promising future. Trends Genet 25, 234–241.

17. Delfarah A, Hartel NG, Zheng D, Yang J & Graham NA (2021) Identification of a Proteomic Signature of Senescence in Primary Human Mammary Epithelial Cells. J Proteome Res 20, 5169–5179.

18. Dimri GP, Lee X, Basile G, Acosta M, Scott G, Roskelley C, Medrano EE, Linskens M, Rubelj I & Pereira-Smith O (1995) A biomarker that identifies senescent human cells in culture and in aging skin in vivo. Proceedings of the National Academy of Sciences 92, 9363–9367.

19. Dodig S, Čepelak I & Pavić I (2019) Hallmarks of senescence and aging. Biochem Med (Zagreb) 29, 030501.

20. Finch CE (2009) Update on slow aging and negligible senescence--a mini-review. Gerontology 55, 307–313.

21. Gomes NMV, Ryder OA, Houck ML, Charter SJ, Walker W, Forsyth NR, Austad SN, Venditti C, Pagel M, Shay JW & Wright WE (2011) Comparative biology of mammalian telomeres: hypotheses on ancestral states and the roles of telomeres in longevity determination. Aging Cell 10, 761–768.

22. Grabowski GA & Mistry PK (2022) Therapies for lysosomal storage diseases: Principles, practice, and prospects for refinements based on evolving science. Mol Genet Metab 137, 81–91.

23. Green DR, Galluzzi L & Kroemer G (2011) Mitochondria and the autophagy-inflammation-cell death axis in organismal aging. Science 333, 1109–1112.

24. Gu Z, Jiang J, Tan W, Xia Y, Cao H, Meng Y, Da Z, Liu H & Cheng C (2013) p53/p21 Pathway involved in mediating cellular senescence of bone marrow-derived mesenchymal stem cells from systemic lupus erythematosus patients. Clin Dev Immunol 2013, 134243.

25. Gupta P, Soyombo AA, Shelton JM, Wilkofsky IG, Wisniewski KE, Richardson JA & Hofmann SL (2003) Disruption of PPT2 in mice causes an unusual lysosomal storage disorder with neurovisceral features. Proceedings of the National Academy of Sciences 100, 12325–12330.

26. Hansen M, Rubinsztein DC & Walker DW (2018) Autophagy as a promoter of longevity: insights from model organisms. Nat Rev Mol Cell Biol 19, 579–593.

27. Harper JM, Salmon AB, Leiser SF, Galecki AT & Miller RA (2007) Skin-derived fibroblasts from long-lived species are resistant to some, but not all, lethal stresses and to the mitochondrial inhibitor rotenone. Aging Cell 6, 1–13.

28. Harr B, Karakoc E, Neme R, Teschke M, Pfeifle C, Pezer Ž, Babiker H, Linnenbrink M, Montero I, Scavetta R, Abai MR, Molins MP, Schlegel M, Ulrich RG, Altmüller J, Franitza M, Büntge A, Künzel S & Tautz D (2016) Genomic resources for wild populations of the house mouse, Mus musculus and its close relative Mus spretus. Sci Data 3, 160075.

29. Hernandez-Segura A, Brandenburg S & Demaria M (2018) Induction and Validation of Cellular Senescence in Primary Human Cells. J Vis Exp, 57782.

30. Hornsby PJ (2002) Cellular senescence and tissue aging in vivo. J Gerontol A Biol Sci Med Sci 57, B251–256.

31. Imai Y, Tanave A, Matsuyama M & Koide T (2022) Efficient genome editing in wild strains of mice using the i-GONAD method. Sci Rep 12, 13821.

32. Ito T, Teo YV, Evans SA, Neretti N & Sedivy JM (2018) Regulation of Cellular Senescence by Polycomb Chromatin Modifiers through Distinct DNA Damage-and Histone Methylation-Dependent Pathways. Cell Rep 22, 3480–3492.

33. Kang T, Moore EC, Kopania EEK, King CD, Schilling B, Campisi J, Good JM & Brem RB (2023) A natural variation-based screen in mouse cells reveals USF2 as a regulator of the DNA damage response and cellular senescence. G3 (Bethesda) 13, jkad091.

34. Kawakami M & Yamamura K-I (2008) Cranial bone morphometric study among mouse strains. BMC Evolutionary Biology, 8 (1), 73.

35. Khan M & Gasser S (2016) Generating Primary Fibroblast Cultures from Mouse Ear and Tail Tissues. J Vis Exp, 53565.

36. Kreis N-N, Friemel A, Zimmer B, Roth S, Rieger MA, Rolle U, Louwen F & Yuan J (2016) Mitotic p21Cip1/CDKN1A is regulated by cyclin-dependent kinase 1 phosphorylation. Oncotarget 7, 50215–50228.

37. Lee BY, Han JA, Im JS, Morrone A, Johung K, Goodwin EC, Kleijer WJ, DiMaio D & Hwang ES (2006) Senescence-associated beta-galactosidase is lysosomal beta-galactosidase. Aging Cell 5, 187–195.

38. Lee J-H, Yu WH, Kumar A, Lee S, Mohan PS, Peterhoff CM, Wolfe DM, Martinez-Vicente M, Massey AC, Sovak G, Uchiyama Y, Westaway D, Cuervo AM & Nixon RA (2010) Lysosomal Proteolysis and Autophagy Require Presenilin 1 and Are Disrupted by Alzheimer-Related PS1 Mutations. Cell 141, 1146–1158.

39. Leiva-Rodríguez T, Romeo-Guitart D, Marmolejo-Martínez-Artesero S, Herrando-Grabulosa M, Bosch A, Forés J & Casas C (2018) ATG5 overexpression is neuroprotective and attenuates cytoskeletal and vesicle-trafficking alterations in axotomized motoneurons. Cell Death Dis 9, 626.

40. Liu X, Ye Q, Huang Z, Li X, Zhang L, Liu X, Wu Y-C, Brockmeier U, Hermann DM, Wang Y- C & Ren L (2023) BAG3 Overexpression Attenuates Ischemic Stroke Injury by Activating Autophagy and Inhibiting Apoptosis. Stroke 54, 2114–2125.

41. Mahieu T, Park JM, Revets H, Pasche B, Lengeling A, Staelens J, Wullaert A, Vanlaere I, Hochepied T, Van Roy F, Karin M, Libert C (2006) The wild-derived inbred mouse strain SPRET/Ei is resistant to LPS and defective in IFN-β production. Proceedings of the National Academy of Sciences 103 (7), 2292–2297.

42. Morgan AP, Hughes JJ, Didion JP, Jolley WJ, Campbell KJ, Threadgill DW, Bonhomme F, Searle JB & de Villena FP-M (2022) Population structure and inbreeding in wild house mice (Mus musculus) at different geographic scales. Heredity (Edinb) 129, 183–194.

43. Oka K, Yamakawa M, Kawamura Y, Kutsukake N & Miura K (2023) The Naked Mole-Rat as a Model for Healthy Aging. Annu Rev Anim Biosci 11, 207–226.

44. Pagès M, Fabre P, Chaval Y, Mortelliti A, Nicolas V, Wells K, Michaux JR & Lazzari V (2015) Molecular phylogeny of South-East Asian arboreal murine rodents. Zoologica Scripta 45 (4), 349–364.

45. Pyo J-O, Yoo S-M, Ahn H-H, Nah J, Hong S-H, Kam T-I, Jung S & Jung Y-K (2013) Overexpression of Atg5 in mice activates autophagy and extends lifespan. Nat Commun 4, 2300.

46. Robbins E, Levine EM & Eagle H (1970) Morphologic changes accompanying senescence of cultured human diploid cells. J Exp Med 131, 1211–1222.

47. Rovira M, Sereda R, Pladevall-Morera D, Ramponi V, Marin I, Maus M, Madrigal-Matute J, Díaz A, García F, Muñoz J, Cuervo AM & Serrano M (2022) The lysosomal proteome of senescent cells contributes to the senescence secretome. Aging Cell 21, e13707.

48. Sami S, Liu G, Hornicek F, Cates JM & Mankin HJ (2002) Membranous lipodystrophy. A case report. J Bone Joint Surg Am 84, 630–633.

49. Saul D, Jurk D, Doolittle ML, Kosinsky RL, Monroe DG, LeBrasseur NK, Robbins PD, Niedernhofer LJ, Khosla S & Passos JF (2023) Distinct secretomes in p16-and p21-positive senescent cells across tissues. bioRxiv, 2023.12.05.569858.

50. Shin HR & Zoncu R (2020) The Lysosome at the Intersection of Cellular Growth and Destruction. Dev Cell 54, 226–238.

51. Shin JY, Hong S-H, Kang B, Minai-Tehrani A & Cho M-H (2013) Overexpression of beclin1 induced autophagy and apoptosis in lungs of K-rasLA1 mice. Lung Cancer 81, 362– 370.

52. Smissen PJ & Rowe KC (2018) Repeated biome transitions in the evolution of Australian rodents. Molecular Phylogenetics and Evolution 128, 182–191.

53. Staelens J, Wielockx B, Puimège L, Van Roy F, Guénet J-L & Libert C (2002) Hyporesponsiveness of SPRET/Ei mice to lethal shock induced by tumor necrosis factor and implications for a TNF-based antitumor therapy. Proceedings of the National Academy of Sciences, 99 (14), 9340–9345.

54. Sulak M, Fong L, Mika K, Chigurupati S, Yon L, Mongan NP, Emes RD & Lynch VJ (2016) TP53 copy number expansion is associated with the evolution of increased body size and an enhanced DNA damage response in elephants. eLife 5, e11994.

55. Tian X, Firsanov D, Zhang Z, Cheng Y, Luo L, Tombline G, Tan R, Simon M, Henderson S, Steffan J, Goldfarb A, Tam J, Zheng K, Cornwell A, Johnson A, Yang J-N, Mao Z, Manta B, Dang W, Zhang Z, Vijg J, Wolfe A, Moody K, Kennedy BK, Bohmann D, Gladyshev VN, Seluanov A & Gorbunova V (2019) SIRT6 Is Responsible for More Efficient DNA Double-Strand Break Repair in Long-Lived Species. Cell 177, 622–638.e22.

56. Wannemacher R, Stegmann F, Eikelberg D, Bühler M, Li D, Kohale SK, Asawapattanakul T, Ebbecke T, Raulf M-K, Baumgärtner W, Lepenies B & Gerhauser I (2025) Infection of a β-galactosidase-deficient mouse strain with Theiler’s murine encephalomyelitis virus reveals limited immunological dysregulations in this lysosomal storage disease. Front Immunol 16, 1467207.

57. Yew T-L, Chiu F-Y, Tsai C-C, Chen H-L, Lee W-P, Chen Y-J, Chang M-C & Hung S-C (2011) Knockdown of p21(Cip1/Waf1) enhances proliferation, the expression of stemness markers, and osteogenic potential in human mesenchymal stem cells. Aging Cell 10, 349–361.

58. Yuan Y, Kilpatrick BS, Gerndt S, Bracher F, Grimm C, Schapira AH & Patel S (2021) The lysosomotrope GPN mobilises Ca2+ from acidic organelles. Journal of Cell Science 134, jcs256578.

59. Zaidi N, Maurer A, Nieke S & Kalbacher H (2008) Cathepsin D: A cellular roadmap. Biochemical and Biophysical Research Communications 376, 5–9.

60. Zhao Y, Tyshkovskiy A, Muñoz-Espín D, Tian X, Serrano M, de Magalhaes JP, Nevo E, Gladyshev VN, Seluanov A & Gorbunova V (2018) Naked mole rats can undergo developmental, oncogene-induced and DNA damage-induced cellular senescence. Proc Natl Acad Sci U S A 115, 1801–1806.

