## Supplemental Materials (Detailed Methods) for "Elevated lysosomal mass and enzyme activity in fibroblasts of the Mediterranean mouse *Mus spretus*"

Supplementary methods text

**Mouse strains and husbandry**

Wild-derived, inbred lines of *Mus musculus musculus* (PWK/PhJ, BLG2/Ms, CHD/Ms,MSM/Ms) and *Mus spretus* (STF/Pas, SFM) were maintained under pathogen-free conditions and controlled lighting (daily light period: 06:00 to 18:00). The mice were housed in a temperature-controlled room (23°C ± 2°C) and food and water were provided ad libitum. The BLG2/Ms (BLG2), CHD/Ms (CHD), and MSM/Ms (MSM) strains, all classified within the *Mus musculus musculus* subspecies group (Imai et al. 2022), were sampled with protocols approved by the Committee for Animal Care and Use of the National Institute of Genetics of Japan (permit number R5-8). The PWK/PhJ, STF/Pas, and SFM strains were sampled with protocols approved by the Institutional Animal Care and Use Committee (IACUC) (IACUC# D16-00210 (A3327-01)). All animals used were aged between 2 and 5 months and were euthanized humanely via CO_2_ treatment for tissue collection. Tails cuttings were collected immediately after sacrifice into chilled fetal bovine serum (FBS) and, if needed, shipped overnight on dry ice for subsequent processing.

The standard laboratory strain *Mus musculus domesticus* (C57BL/6J) (IACUC# A4107-01), wild-derived inbred lines of *Mus musculus domesticus* (ManB/NachJ, TUCA/NachJ) (IACUC# AUP-2017-08-10248), and *Mus spicilegus* (ZRU) (IACUC# AUP-2015-11-8145) were maintained under specific pathogen-free conditions with controlled lighting (12-hour to 14-hour light cycle; lights on from 06:00 to 20:00). The mice were housed in a temperature-controlled room (21 - 22°C) and food and water were provided ad libitum. All animals used in this study were between 3 and 5 months old and were euthanized humanely via isoflurane anesthesia followed by cervical dislocation to collect tissues. Tail cuttings were collected immediately post-sacrifice and placed individually in separate 15mL conical tubes for tissue collection.

**Primary cell extraction and culturing**

We extracted primary tail fibroblasts from the cuttings essentially as described (Khan and Gasser 2016). Briefly, we soaked the tail cuttings in 70% ethanol for 5 minutes and then laid them out in sterile 10 cm tissue culture dishes in a biosafety cabinet for 5 minutes to dry. Next, we transferred the tails to a 10-cm dish containing 10 mL of complete medium composed of RPMI 1640 (Corning cat. #17105CV), 10% fetal bovine serum (Genesee Scientific cat. #25-514H), 50µM 2-mercaptoethanol (MP Biomedicals cat. #194705), 100µM asparagine (Sigma-Aldrich cat.#C980A53), 2mM glutamine (Sigma-Aldrich cat. #G8540), and 1% penicillin-streptomycin (pen-strep) (Corning cat. #30002CI). We removed hair from the tails using sterile forceps and a razor blade, and then cut the tails into 2-3 mm pieces using sterile surgical scissors. These pieces were transferred into a 2 mL cryotube containing a collagenase D–pronase (Thomas Scientific cat. #C756Y81, VWR cat. #97062-916) solution and left in a shaking incubator at 37˚C for 90 minutes. After incubation, we placed the contents of the cryotubes into a 70 µm cell strainer in a new 10-cm dish containing 10 mL of complete medium. Using the back end of a sterile 10 mL syringe plunger, we ground the tissue into the strainer for 5-10 minutes to release the cells into the media. The cell suspension was then collected into 15 mL conical tubes and centrifuged at 580 x g and 4˚C for 5 minutes. We removed the supernatant and replaced it with fresh complete medium, repeating the centrifugation for two additional rounds. After the final spin, we replaced the media with complete medium supplemented with 250 ng/mL of amphotericin B (Thomas Scientific cat. #CHM00A358). The cell suspension was placed in a T25 flask and incubated in a 37˚C humidified incubator at 3% O_2_ and 10% CO_2_ for two days before passage and continued culture in complete medium (DMEM, 10% FBS, 1% pen-strep).

For long term storage, we suspended cells in 5% DMSO (Sigma-Aldrich cat. #D2438) in FBS, aliquoted into cryotubes, and placed the vials in a slow cooling container in a -80˚C freezer overnight. The following day, we moved the cryotubes out of the slow cooling container and placed them into long term containers in the freezer. To thaw cells for continued culture, we swirled cryotubes containing frozen aliquots in a 37˚C water bath until only a thin layer of ice remained in the vial. We then immediately transferred contents of the cryotube into a flask containing pre-warmed complete medium and incubated the flask overnight. The next day, we washed the cells with two washes of PBS to thoroughly remove any residual DMSO before we replenished the flask with complete medium for subsequent culture.

For cell passaging, cells were rinsed with PBS, followed by the addition of trypsin-EDTA 0.25% (ThermoFisher Scientific cat. #25200056), which was incubated with the culture for 2 minutes. The flask or plate was gently tapped to aid in cell detachment. Once detached, trypsin was neutralized with an equal volume of working media, and the cells were split after confirming complete detachment from the plate bottom.

**Irradiation treatment**

The day before irradiation, we seeded approximately 10,000 cells in each of the 6 wells to achieve 60-70% confluency the next day. The plates were then placed in a 37˚C humidified incubator with 3% O_2_ and 10% CO_2_ overnight in complete medium. The day after seeding, the culture was transferred into an X-RAD 320 X-Ray Biological Irradiator and treated with 15 Gy of X-ray irradiation for all experiments except the dose-response, which used 10 Gy, 15 Gy, 20 Gy, and 25 Gy. After irradiation treatment, we replaced the complete medium in the cultures and placed it back into the 37˚C humidified incubator at 3% O_2_ and 10% CO_2_, incubating as appropriate with changes of complete medium every 4 to 5 days.

**Neocarzinostatin treatment**

The day prior to neocarzinostatin treatment, we seeded approximately 10,000 cells in each of the 6 wells to achieve 60-70% confluency the next day. The plates were then placed in a 37˚C humidified incubator with 3% O_2_ and 10% CO_2_ overnight in complete medium. The day after seeding, we mixed neocarzinostatin (Sigma-Aldrich cat. #N9162) with complete medium to achieve a final concentration of 3.6 µM for the treatment solution, making sure to have at least 2mL of treatment solution per replicate well. We then removed the complete medium from the plate, and added the treatment solution to incubate in a 37˚C humidified incubator for 1 hour. Following treatment, cells were washed with two washes of PBS before replenishing with complete medium and returning the cultures into the 37˚C humidified incubator at 3% O_2_ and 10% CO_2_. Any subsequent assays were conducted after a minimum recovery period of 24 hours.

**Senescence assay**

For a given replicate of a fibroblast culture treated with irradiation or neocarzinostatin as above, we measured β-galactosidase (β-gal) activity using the Abcam Ltd. Senescence Detection Kit (Cat. #ab65351). To stain for β-gal activity, the cells were fixed and permeabilized at room temperature before incubated overnight with a staining solution containing XGal overnight. Using a brightfield microscope, we captured images of the stained cells the next day for subsequent manual counting. We repeated this procedure, from seeding to staining, for at least two batches of each genotype and treatment across at least two separate days. In Figure S2, β-gal staining for each species after treatment was normalized to the average corresponding untreated control β-gal staining measure for that species. In Figure 1, we carried out Wilcoxon tests to compare β-gal staining between *M. spretus* strains (STF/Pas, SFM) and that of all other genotypes under each condition in turn (control, irradiated, and neocarzinostatin). We also carried out Wilcoxon tests comparing *M. spretus* strains (STF/Pas, SFM) to *M. spicilegus* under each condition in turn, and we did a two-factor ANOVA with treatment and genotype as factors. For Figure S2, we used β-gal staining as input into a Wilcoxon test comparing *M. spretus* strains (STF/Pas, SFM) to all other genotypes for each normalized data set in turn (irradiation normalized to control and, separately, neocarzinostatin-treated normalized to control). For Figure S5, we used β-gal staining as input into a two-factor ANOVA with genotype and irradiation dose as factors.

**Apoptosis marker assay**

We used the ApoTox-Glo™ Triplex Assay (Promega cat#G6321) to assess apoptosis. For both neocarzinostatin and irradiation treatment, we seeded approximately 5,000 cells in each of the 96 wells to achieve 60-70% confluency the day of treatments. The plates were then placed in a 37˚C humidified incubator with 3% O_2_ and 10% CO_2_ overnight in complete medium for treatment the next day. We repeated this procedure, from seeding to staining, for at least two batches of each genotype and treatment across at least two separate days. After irradiation or neocarzinostatin treatment as above, a given replicate was analyzed in a SpectraMax M3 Microplate Reader. Untreated control replicates were plated the day before staining and measurement.

**LysoTracker staining, flow cytometry, and microscopy**

We used LysoTracker™ Green DND-26 (Invitrogen) to assess lysosomal titer via quantification of intracellular acidity as follows. For both neocarzinostatin and irradiation treatment, we seeded approximately 10,000 cells in each of 6 wells to achieve 60-70% confluency the day of treatments. The plates were then placed in a 37˚C humidified incubator with 3% O_2_ and 10% CO_2_ overnight in complete medium for treatment the next day. After irradiation or neocarzinostatin treatment as above, the cells were incubated in 2 mL media containing 0.024% LysoTracker for 1 hour, then washed with PBS. For staining, cells were incubated for 1 hour in culture media containing LysoTracker at a working concentration of 50 nM at 37°C  prior to imaging. Staining of a given replicate was quantified in an Attune Flow Cytometer or Zeiss LSM880 FCS confocal microscope with the absorption and emission set at 504 and 511 nm, respectively, and gain kept constant. For each treatment group (control, neocarzinostatin, and irradiation), at least 10,000 cells were quantified by flow cytometry. We repeated this procedure, from seeding to staining, for at least two batches of each genotype and treatment across at least two separate days. Untreated control replicates were plated the day before staining and measurement. For microscopy-based quantification, images were obtained using Zen (black edition) software and quantified using ImageJ. The images were binarized, converted to 8-bit, and thresholded using the IsoData auto-thresholding method. We used as a readout the number of fluorescent pixels per cell. For Figure 2A and S9, we carried out a Wilcoxon test comparing *M. spretus* (STF/Pas) to *M. m. musculus* (PWK/PhJ) under each condition in turn (control, irradiated, and neocarzinostatin).

**BODIPY FL pepstatin A assay and microscopy**

We used staining of BODIPY-FL pepstatin A (Invitrogen) to assess cathepsin D activity as follows. For both neocarzinostatin and irradiation treatment, we seeded approximately 5,000 cells in each of 12 wells to achieve 60-70% confluency the day of treatment. For staining, cells were incubated for 30 minutes in culture media containing BODIPY FL pepstatin A at a working concentration of 70 nM at 37°C prior to imaging. Image acquisition and processing was as for Lysotracker, above, except staining of a given replicate was quantified with Zeiss LSM880 FCS confocal microscope and Zen (black edition) software, with the absorption and emission set at 590 and 645 nm, respectively. We repeated this procedure, from seeding to staining, for at least two batches of each genotype and treatment across at least two separate days. Untreated control replicates were plated the day before staining and measurement, microscopy-based quantification was performed with the procedure previously described. For Figure 2B, we carried out a Wilcoxon test comparing *M. spretus* (STF/Pas) to *M. m. musculus* (PWK/PhJ) under each condition in turn (control, irradiated, and neocarzinostatin).

**P21 Western blotting**

For a given replicate of P21 protein quantification by Western blot, after irradiation or neocarzinostatin treatment as above, we harvested approximately 1 million primary fibroblast cells, using trypsin to detach the cells; untreated control replicates were plated the day before measurement. Collected cells were washed with PBS and centrifuged at 1000 x g for 5 minutes. After aspirating the supernatant, the cell pellet was prepared for total protein extraction in a buffer containing 1% SDS, 1% NP-40, 150 mM NaCl, 50 mM Tris-HCl (pH 8), and 0.5% sodium deoxycholate, along with Halt Protease and Phosphatase Inhibitor Cocktail (Thermo Scientific). Lysates were sonicated (qSonica Ultrasonicator, 5 min at 75A) and protein concentrations were measured using a Bradford protein assay (Bio-Rad). Equal amounts of whole cell lysate protein (25 µg) were denatured in Laemmli Buffer (Bio-Rad), resolved by SDS-PAGE, and transferred to a PVDF membrane (Bio-Rad) using the Trans-Blot Turbo Transfer System (Bio-Rad, Mixed Molecular Weight protocol). Western blotting was performed in Tris-buffered saline with 0.2% Tween-20 (TBS-T), with PageRuler Prestained Protein Ladder (10–180 kDa; ThermoFisher) for molecular weight reference. Primary antibodies included rabbit monoclonal Anti-p21 (ab188224, Abcam, 1:1000) and mouse monoclonal anti-β-tubulin (T9026, Sigma-Aldrich, 1:1000). Secondary antibodies used were Goat Anti-Mouse IgG(H+L) Human ads-HRP (Cat#1031-05, Southern Biotech, 1:5000) and Goat Anti-Rabbit IgG(H+L) Human ads-HRP (Cat#4050-05, Southern Biotech, 1:5000). Blots were developed using either Clarity Western ECL Substrate (Bio-Rad) or Radiance Plus Femtogram-sensitivity HRP Substrate (Azure Biosystems) and imaged on a Chemidoc MP imager (Bio-Rad). ImageJ was used for preparing blots for publication. At least three biological replicates of a given genotype in a given condition were carried out across at least three separate days.

For Western blot quantification, ImageJ was used to define the region of interest (ROI) for each scanned blot, with the largest protein band in the row serving as the standardized size per blot. The mean gray value of the selected area was measured for each sample. Background intensity was corrected by subtracting the background gray value from the ROI measurement for each sample. This process was repeated for the loading control (tubulin) and its corresponding background. The final quantification was calculated as the ratio of the net protein signal to the net loading control signal. For Figure S7, we used the resulting values as input into a two-factor ANOVA with conditions and species as factors.

**Bafilomycin A1 treatment**

We used Bafilomycin A1 (Cell Signaling Technology) as a negative control for LysoTracker staining, to disrupt the function of the lysosomal H+-ATPase, as follows. We seeded approximately 5,000 cells in each of 12 wells to achieve 60-70% confluency the day of treatments. The plates were then placed in a 37˚C humidified incubator with 3% O_2_ and 10% CO_2_ overnight in complete medium. The following day, the cells were incubated with 100 nM of bafilomycin A1 in media for 30 minutes at 37°C with 3% O_2_ and 10% CO_2_. LysoTracker staining was performed immediately in media containing 100 nM bafilomycin A1, following the staining protocol described above. Cells were incubated under the previously described conditions for 1 hour, after which the staining solution was removed and replaced with 100 nM bafilomycin A1 in PBS for imaging. Fluorescence from each replicate was imaged using a Zeiss LSM880 FCS confocal microscope and Zen (black edition) software, with excitation and emission wavelengths set to 504 nm and 511 nm, respectively, and gain kept constant. Fluorescence was quantified with ImageJ and the same software settings previously described.

We used Alamar Blue (Bio-Rad, cat#BUF012A) to confirm that bafilomycin A1 treatment did not affect cell viability. Approximately 3,000 cells were seeded per well in a 96-well plate one day prior to treatment to ensure consistent confluency. Cells were treated with bafilomycin A1 as described above, except the incubation period was 1 hour and 30 minutes. The medium was then replaced with 100 μL of fresh medium containing 10 μL (1/10th volume) of Alamar Blue, following the manufacturer’s protocol. After 72 hours of incubation at 37°C with 3% O₂ and 10% CO₂, fluorescence was measured using an Infinite® 200 PRO Tecan plate reader at 570 nm to assess cell viability.

**Gene expression analysis**

For Figures S6, S8, and S11, we obtained normalized RNA-sequencing expression data from individual replicates of primary fibroblasts from *M. m. musculus* PWK, *M. spretus* STF, and their F1 hybrid under untreated control and irradiation-treated conditions from Kang et al. (2023) and can be found in the NCBI Gene Expression Omnibus (GEO; https://www.ncbi.nlm.nih.gov/geo/) under accession number GSE201217. For analyses presented in Figures S6 and S8, we applied a two-factor ANOVA with treatment condition and genotype as independent variables.
